# Transcriptional response of the xerotolerant *Arthrobacter* sp. Helios strain to PEG-induced drought stress

**DOI:** 10.1101/2022.07.26.501551

**Authors:** G Hernández-Fernández, B Galán, M Carmona, L Castro, JL García

## Abstract

A new bacterial strain highly tolerant to desiccation and to UV radiation has been isolated from the microbiome of solar panels. This strain showed a high xerotolerance in the exponential and the stationary phase of growth and it has been classified as *Arthrobacter* sp. Helios according to its 16S rDNA, positioning this new strain in the ‘*Arthrobacter citreus* group’. The complete genome of *Arthrobacter* sp. Helios consists in a single circular chromosome of 3,895,998 bp, with a 66% GC content and no plasmids. A total of 3,586 genes were predicted, of which 2,275 protein-encoding genes were functionally assigned. The genome analysis suggests that it is motile, ecologically versatile, capable of growing in a variety of carbon sources and well poised to respond to environmental stresses. Using PEG6000 to mimic arid stress conditions, we have studied the transcriptional response of this strain to matric stress when cells are cultured on media containing 10% (PEG10) and 35% PEG (PEG35). The transcriptomic analysis revealed that cells can be easily adapted to moderate matric stress (PEG10) by modifying the expression of a small number of genes to maintain a high growth rate, while a higher matric stress (PEG35) altered the expression of many more genes. Remarkably, these metabolic changes do not confer the cells a higher tolerance to desiccation, suggesting that mechanisms to support matric stress and desiccation tolerance are different. The peculiar observation that *Arthrobacter* sp. Helios seems to be permanently prepared to handle the desiccation stress makes it an exciting chassis for biotechnological applications.

## 1. Introduction

Bacteria of the genus *Arthrobacter*, belonging to *Micrococcaceae* family and *Actinobacteria* phylum, are among the most frequently isolated, indigenous, aerobic bacterial genera found in soils. Members of the genus are metabolically and ecologically diverse and have the ability to survive in environmentally harsh conditions for extended periods of time (Conn and Dimmick, 1947; Koch et al., 1995; Unell et al., 2008). They have been isolated worldwide from a variety of environments, including sediments (Dastager et al., 2014), human clinical specimens (Huang et al., 2005), water (Kim and Quan, 2007), glacier cryoconite (Margesin et al., 2012), sewage (Kim et al., 2008), glacier ice (Liu et al., 2018) and even from contaminated environments with industrial chemicals and radioactive materials (Sánchez-Castro et al., 2017). The prevalence of *Arthrobacter* in soils can be due to its metabolic versatility and to its ability to survive long periods under stressful conditions such as starvation, temperature shifts, dryness, ionizing radicals and metals, among others (Mongodin et al., 2006; Yao et al., 2015; Porcar et al., 2018). *Arthrobacter* are described to grow as two distinct shapes, forming either spherical or rod-shape cells depending upon culture medium and growth phase, but both forms appear to be equally resistant to desiccation and starvation (Jones and Keddie, 2006; Santacruz-Calvo et al., 2013).

Arid stress caused by the lack of water or by high concentration of salts in the environment is one of the most common stresses that bacteria face in the natural environment (Holden et al., 1997; Holden and Fierer, 2005). Surface soils are unsaturated habitats where the water fluctuation is one of the major factors affecting bacterial cells. Xerotolerant strains tolerate low water activities (*a*_w_) caused either by an increased extracellular osmolarity or by a general lack of water. Osmotic stress is a constant challenge for bacteria living in a range of soils that also affects symbionts that must alternate between two main biotopes, the soil and the root tissues (Hawkins and Oresnik, 2022). In non-saline soils, capillary forces and physical sorption of water to solids, together constituting the soil matric potential, are the dominating factors determining water availability (Holden, 2001). Low matric potentials (i.e., desiccation) limit transport and diffusion of nutrients, impair microbial mobility, and negatively affect the physiological activity of soil bacteria (Stark and Firestone, 1995; Or et al., 2007; Dechesne et al., 2008). Two strategies have evolved to allow microbes to counterbalance the extracellular osmolarity. The first one is the accumulation in the cytoplasm of K^+^ and their counterion glutamate to provide osmotic balance to the cells (Galinski and Trüper, 1994; Epstein, 2003; Gunde-Cimerman et al., 2018). The second strategy consists in the accumulation of small organic compounds, either by *de novo* synthesis or by their uptake from the environment (Oren, 1999; Gunde-Cimerman et al., 2018). These compounds are named compatible solutes because they can be accumulated in high concentrations while not interfering with the metabolism. Most common compatible solutes are sugars (e.g., trehalose), polyols (e.g., mannitol), amino acids (e.g., glutamate) and derivatives thereof (e.g., ectoine) as well as trimethylammonium compounds (e.g., glycine betaine) (Potts 1994; Ruhal et al. 2013; Lebre et al. 2017; Zeidler and Müller 2019).

Remarkably, solar panels provide a new non-natural extreme environment to the microorganisms attached to them, since they are exposed to high and cyclic variations of temperature, sunlight, radiation and humidity, thus resulting in a particular harsh habitat where only heat-, desiccation- and irradiation-adapted microorganisms would survive. Moreover, the surface of solar panels represents a harsh environment exposed to lack of nutrients and water (Dorado-Morales et al., 2016; Porcar et al., 2018). However, solar panels harbour a highly diverse microbial community, including more than 500 different species per panel, most of which belong to drought, heat and radiation-adapted bacterial genera (Dorado-Morales et al., 2016; Tanner et al., 2018; Castillo et al., 2021).

In this work, we have isolated from a solar panel the bacterium *Arthrobacter* sp. Helios, that showed several extremophile properties, but especially a high xerotolerance. Its genome sequence and several physiological characteristics are described. The transcriptome in the presence of PEG6000 simulating matric stress has been analysed to determine the strategies developed by this bacterium to resist water stress. The biotechnological applications of this new strain are discussed.

## 2. Materials and Methods

### 2.1. Isolation of xerotolerant strains from solar panels

Sampling of solar panels was performed as described (Dorado-Morales et al., 2016). Briefly, harvesting of microbiota was carried out by pouring sterile phosphate-buffered saline (PBS) (pH 7.4) on the solar panel and scraping the surface with a window cleaner attached to an autoclaved silicone tube (5 mm in diameter). The resulting liquid suspension was collected by using a pipette and transferred to Falcon tubes, placed on ice, and immediately transported to the lab, where it was filtered by a hydrophilic nylon membrane (Millipore, Burlington, MA, USA) (20 μm pore size) to discard particles, most of the fungi, and inorganic debris. A number of 100 μL aliquots of microbiota samples resuspended in saline solution 0.85% (w/v) obtained from solar panels were spread into Millipore™ membrane filters (0.45 μm pore size, 47 mm diameter, mixed cellulose esters, hydrophilic), air dried and incubated in a stove at 37 °C with 10–15% humidity for 10 days. Filters were hydrated with 1 mL of PBS and the bacterial suspension was plated on LB agar and incubated overnight at 37 °C. The isolated colonies were cultured individually in liquid LB medium and subjected to an additional desiccation cycle in order to obtain the most xerotolerant strains. The validation of the xerotolerance test was performed using *Escherichia coli* DH10B, as a bacterial reference for low xerotolerance; and *Deinococcus radiodurans*, for high xerotolerance. To carry out the comparative tests, 100 μL of the strain cultures in rich media with an optical density at 600 nm (*OD_600_*) of 0.05 were deposited on the filters and incubated in a stove at 37 °C with 10–15% humidity for several days. The filters were hydrated with 1 mL of PBS after 3, 7, and 15 days. To quantify the survival ratio, viability cell count was performed on LB agar plates. The identification of the xerotolerant isolated strains was made by 16S rRNA sequencing. A 1340 bp conserved fragment of 16S rRNA gene was amplified by PCR from genomic DNA, using universal primers 63F (5′-CAGGCCTAACACATGCAAGTC-3′) and 1387R (5′-GGGCGGWGTGTACAAGGC-3′) (Marchesi et al., 1998). PCR products were checked in 0.7 % agarose gel and purified with QIAquick PCR Purification Kit. Sequencing was carried out by Secugen S.L. (Madrid, Spain). The resulting sequences were compared to the nucleotide collection at NCBI using the BLAST tool (http://blast.ncbi.nlm.nih.gov/Blast.cgi) optimized for highly similar sequences (megablast).

### 2.2. Culture and growth and other extremophilic traits

The bacterial strains used in this work were *Arthrobacter* sp. Helios, *Arthrobacter koreensis* CA15-8, *Exiguobacterium* sp. Helios, *Pseudomonas putida* KT2440, *Escherichia coli* DH10B and *Deinococcus radiodurans. Arthrobacter* sp. Helios, *A. koreensis* CA15-8, *Exiguobacterium* sp. Helios and *P. putida* KT2440 were grown in LB medium at 30 °C with orbital shaking at 200 rpm, while *E. coli* DH10B was grown at 37 °C in the same conditions. *D. radiodurans* was grown at 30 °C and 200 rpm in TGY medium (tryptone 1%, glucose 0.1% and yeast extract 0.5%, pH 7.2).

Minimal media M63 supplemented with trace elements and vitamins (Cohen and Rickenberg 1956) was used to study the ability of *Arthrobacter* sp. Helios to grow in different carbon sources as a sole source of carbon and energy. The carbon sources tested were 10 mM glucose, 3 mM fructose, 10 mM xylose, 17 mM succinate, 10 mM maltose, 10 mM sucrose, 10 mM galactose, 8 mM citrate, 10 mM lactose, 10 mM arabinose, 3 mM ribose, 1 mM phenol, 3 mM benzoic acid, 3 mM 3-hydroxybenzoic acid, 3 mM 4-hydroxybenzoic acid, 3 mM protocatechuate, 3 mM catechol, 3 mM phenylacetic acid, 1 mM cholesterol, 3 mM pyridine, 3 mM phthalate, 3 mM isophthalate and 3 mM terephthalate. All products were purchased from Merck and bacterial growth was assessed measuring culture turbidity (*OD_600_*).

For the salinity tolerance test, cells were grown in LB medium supplemented with increasing concentrations of NaCl (20-100 g/L) and *OD_600_* was monitored. In the case of strains UV resistance, cells were grown until the stationary phase, washed twice with PBS and *OD_600_* adjusted to 0.5. 2 mL of each strain were spread in MW6 (Falcon) and serial dilutions were made in order to know the initial CFU/mL. A UV Stratalinker 1800 (Cultek) was used to irradiate cells with increasing doses of UV. Finally, after the exposure, serial dilutions were plated on LB agar plates and colonies were grown overnight to quantify cell viability.

*Arthrobacter* sp. Helios metals and metalloids resistance was assessed in Tris-Minimal Medium (6.06 g/L Tris-HCl; 4.68 g/L NaCl; 1.49 g/L KCl; 1.07 g/L NH_4_Cl; 0.43 g/L Na_2_SO_4_; 0.2 g/L MgCl_2_ 6H_2_O; 0.03 g/L Ca_2_Cl 2H_2_O; 0.23 g/L Na_2_HPO_4_ 12 H_2_O; 0.005 g/L Fe(III)NH4 citrate; 1 μL/L 25% HCl; 70 μg/mL ZnCl_2_; 100 μg/mL MnCl_2_ 4H_2_O; 60 μg/mL H_3_BO_3_; 200 μg/mL CoCl_2_ 6H_2_O; 20 μg/mL CuCl_2_ 2H_2_O; 20 μg/mL NiCl_2_ 6H_2_O; 40 μg/mL Na_2_MoO_4_ 2H_2_O) supplemented with yeast extract (1 g/L). The metals and metalloids were added to this medium in the following concentrations: 0.62-1.25 mM NiCl_2_, 0.62-10 mM ZnCl_2_ and K_2_TeO_3_, 0.62 mM NaAsO_2_, 1 mM Na_2_HAsO_4_, 0.4-3 mM CuSO_4_, 0.62 mM CdCl_2_ and AgNO_3_ and 0.62-2.5 mM Pb(NO_3_)_2_. *Arthrobacter* sp. Helios selenite resistance was tested in LB medium with 1-200 mM Na_2_SeO_3_ and *OD_600_* monitored. For the SeNPs production, *Arthrobacter* sp. Helios was grown in LB medium supplemented with 1 mM Na_2_SeO_3_ for 24 h at 30 °C with orbital shaking at 200 rpm.

To test the ability of *Arthrobacter* sp. Helios to grow in PEG-mediated drought stress, polyethylene glycol 6000 (PEG6000) (Sigma-Aldrich) was used. PEG6000 was added to LB medium in concentrations of 10, 20, 30 and 35% (w/v) to assess *Arthrobacter* sp. Helios resistance. Concentrations of 10% and 35% were selected for transcriptomic analysis. To inoculate 10% and 20% PEG6000 cultures, bacteria were previously grown overnight in LB liquid medium without PEG6000, whereas 30% and 35% PEG6000 cultures were inoculated with *Arthrobacter* sp. Helios previously grown overnight in the presence of 20% PEG6000. All of these cultures started at an initial *OD_600_* of 0.1.

### 2.3. Characterization of the Selenium Nanoparticles (SeNPs) and Transmission Electron Microscopy (TEM)

Samples of *Arthrobacter* sp. Helios grown with 1 mM sodium selenite were dropped onto carbon-coated copper grids allowing the solvent to evaporate. TEM analyses were performed with a JEOL model JEM-2100 instrument operated at an accelerating voltage of 200 kV. The elemental composition of the selenium nanoparticles was determined by energy-dispersive X-ray spectroscopy (EDX) (Li et al. 2014; Collins 2007). SeNPs size distribution was calculated using the Image J software (https://imagej.nih.gov/ij/download.html).

### 2.4. Sequencing, Assembly, and Bioinformatic Analyses of Genome

Total DNA extraction and sequencing was performed as described (Castillo et al., 2021). Briefly, *Arthrobacter* sp. Helios genome was sequenced and assembled by Microbes NG (https://microbesng.com/) using Illumina technology and its standard pipeline for a *de novo* assembly. Kraken (https://ccb.jhu.edu/software/kraken/) was used to identify the closest available reference genome and reads were mapped to this reference genome using BWA mem (Burrows–Wheeler Aligner, http://bio-bwa.sourceforge.net/) in order to check the sequencing data quality. A *d*e *novo* assembly of the reads was performed using SPAdes, and reads were mapped back to the resultant contigs, using BWA mem to obtain more quality metrics. The number of reads was 1,483,364 with a median insert size of 156 bp and a mean coverage of 147. The number of contigs delivered was 45 with a N50 of 147,883, being the largest contig size of 317,147 bp. To enhance genome quality, we further sequenced the genome with Nanopore technology. A genomic library was created with the 1D Native Barcoding genomic DNA Barcode kit and run through the flow cell FLO-MIN-106D v R9 in a MinION equipment. The number of reads obtained was 406,031 with a median insert size of 2,480 and an average quality of 11.32.

The assembly was performed using the Galaxy Community Hub (https://galaxyproject.org/), first selecting the reads longer than 1 kb and with a quality bigger than 10 using Filtlong software (v 0.2.0) (https://github.com/rrwick/Filtlong) and comparing them with the Illumina reads using Unicycler (v 0.4.8) (https://github.com/rrwick/Unicycler/releases/tag/v0.4.8) with the standard parameters. This resulted in one single contig of 3,895,998. The genome was structurally annotated using the RAST Server (https://rast.nmpdr.org/), and automated genome annotation system, functions, names, and general properties of gene products were predicted using this method. Phylogenetic analyses were performed using Geneious v *2022.0.1* software (http://www.geneious.com). Comparative analyses were carried out using BLAST software at NCBI (https://blast.ncbi.nlm.nih.gov/Blast.cgi). The genome project has been deposited at GenBank under the accession number CP095402.

### 2.5. Biomass collection and RNA extraction

Total RNA was extracted from cultures of *Arthrobacter* sp. Helios grown in 50 ml LB medium supplemented with 10% (PEG10) and 35% (PEG35) PEG6000, as the simulated drought conditions, and 0% (PEG0) PEG6000 as the control condition. Bacteria were grown until the middle of their exponential growth and biomass was collected as follows: 10 mL of each culture were centrifuged at 4 °C for 10 min at 3800 rpm in an Eppendorf Centrifuge 5810 R and then washed twice with PBS. 1 mL of a solution containing SDS 1% (v/v), 160 mM EDTA and lysozyme (50 mg/mL) was added to the cell pellet and then transferred to a 15 mL Falcon tube with glass balls previously sterilized. The solution was left at RT for 5 min in order to let the lysozyme lyse the cells. Next, 200 μL of phenol-chloroform-isoamyl alcohol (ROTH) were added and vortexed to obtain the cell lysate. Then, 800 μL of buffer RLT (Qiagen) with β-mercaptoethanol (100:1) were added and incubated on ice during 10 min. 3 cycles of vortex followed by an incubation in ice were performed, and 1.4 mL of phenol-chloroform isoamyl alcohol added. The Falcon tubes were centrifuged 15 min and the aqueous phase transferred to a new tube with 700 μL of ethanol. Finally, the total RNA was purified from this extraction using a RNeasy kit (Qiagen) following the manufacturer's instructions.

### 2.6. Transcriptomic analysis

Three biological replicates of each condition were used to sequence the total RNA of *Arthrobacter* sp. Helios grown in PEG-mediated drought stress. The *de novo* transcriptome sequencing was performed by Macrogen NGS Service (https://dna.macrogen.com/) (Illumina TruSeq RNA library, 6 GB/sample sequencing coverage) and fragments of 151 bp paired-end reads were obtained. Raw reads were trimmed and cleaned with Trimmomatic 0.39 in order to remove Illumina adapters and low-quality bases (Bolger et al., 2014). After filtration, we obtained more than 58,000,000 reads for each sample and a mapping ratio against *Arthrobacter* sp. Helios genome ranging between 90-98% (table S1). Besides, the quality score Q30 was above 95%. Trimmed reads were aligned to *Arthrobacter* sp. Helios genome (accession number CP095402) using Bowtie2 2.4.2 (Langmead and Salzberg 2012), and reads were counted with Htseq-count 0.13.5 (Anders et al., 2015). Differential gene expression analysis between groups was performed with the DESeq2 1.32.0 from the R software 3.6.3 (R: The R Project for Statistical Computing. https://www.r-project.org/). Genes with a |log_2_FC|≥_2_ and FDR<0.05 (FC: Fold change; FDR: False discovery rate) were considered as differentially expressed. A Principal Component Analysis (PCA) with the variance stabilizing transformed counts helped to inferred the general quality of the experimental design, i.e., the absence of covariates or batch effects. Heatmap, volcano plots and Venn diagrams were drawn with “ComplexHeatmap”, “VennDiagram” and “EnhancedVolcano”, respectively, from the R package.

The eggNOG-mapper tool (http://eggnog-mapper.embl.de/) (Cantalapiedra et al., 2021) was used for the functional annotation of the proteins with COG and KEGG databases (https://www.ncbi.nlm.nih.gov/research/cog/ and https://www.genome.jp/kegg/, respectively). An enrichment analysis was performed with the function enricher of the “clusterProfiler” R package in order to identify if either up or down regulated genes were significantly overrepresented in each COG category through a hypergeometric test (*=p.adjust<0.05; **=p.adjust<0.01).

## 3. Results

### 3.1. Isolation and identification of the xerotolerant strain Arthrobacter sp. Helios

*Arthrobacter* sp. Helios was isolated from a solar panel due to its high resistance to desiccation since it survived the xerotolerance test described in the methods section. It was identified as *Arthrobacter* sp. according to its 16S rDNA. A BLAST analysis of the 16S rRNA gene sequence from *Arthrobacter* sp. Helios revealed a high similarity with *Arthrobacter luteolus* CF-25 (DSM 13067) and *Arthrobacter koreensis* CA15-8 (DSM 16760), positioning this new strain in the ‘*Arthrobacter citreus* group’ proposed by Busse (2016). *Arthrobacter* sp. Helios showed different morphology (coccoid or rod-shape) depending on the growth phase (Fig. 1).

**Figure 1.**
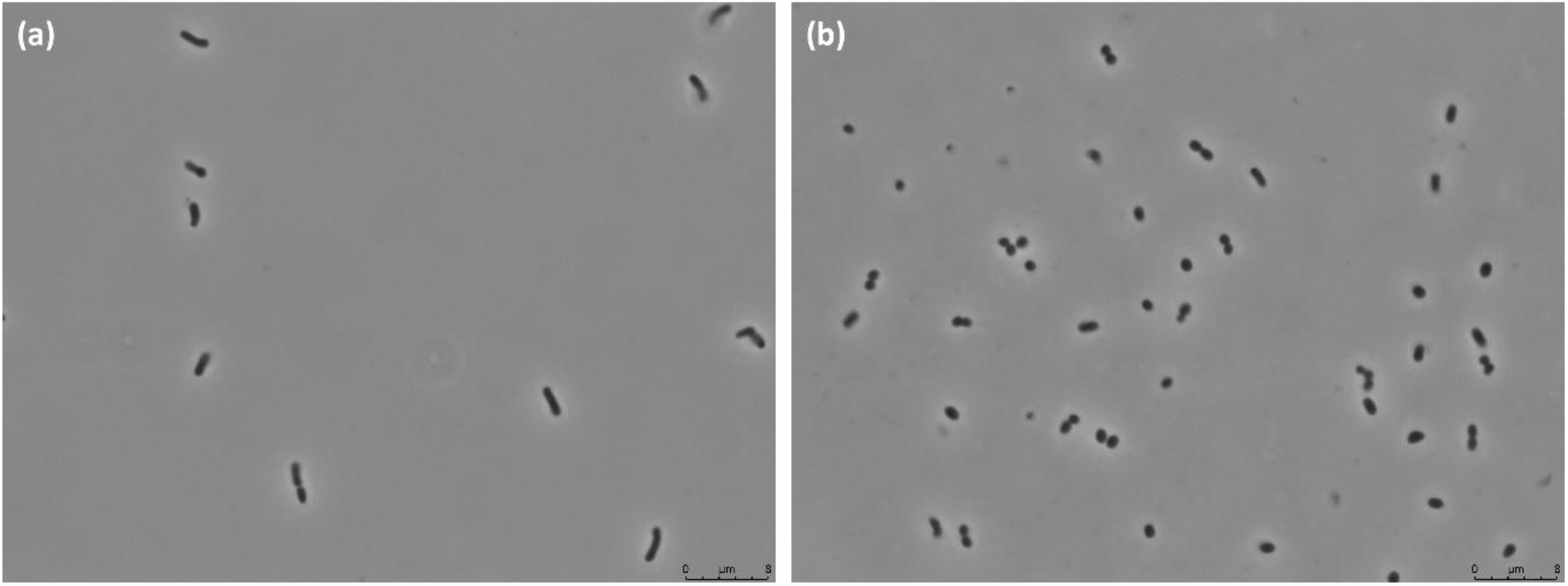
Phase contrast microscopy of *Arthrobacter* sp. Helios grown in LB at (a) exponential phase and (b) stationary phase.

To compare the desiccation resistance of *Arthrobacter*. sp. Helios with other known xerotolerant strains, we performed a desiccation test using *D. radiodurans, Exiguobacterium* sp. Helios (Castillo et al., 2021), the phylogenetically related strain *A. koreensis* CA15-8 (Lee et al., 2003), and *E. coli* DH10B as a negative control. As expected, approximately an 80% of *D. radiodurans* cells survived after desiccation and no survival of *E. coli* cells were detected at any studied time (Fig. 2). Xerotolerance of *Exiguobacterium* sp. Helios and *A. koreensis* was lower than that of *D. radiodurans* with a viability of less than 10% after 3 days in the tested conditions (Fig. 2). However, the survival rate for *Arthrobacter*. sp. Helios was around 30% suggesting a better xerotolerance capacity than the related xerotolerant strain *A. koreensis*.

**Figure 2.**
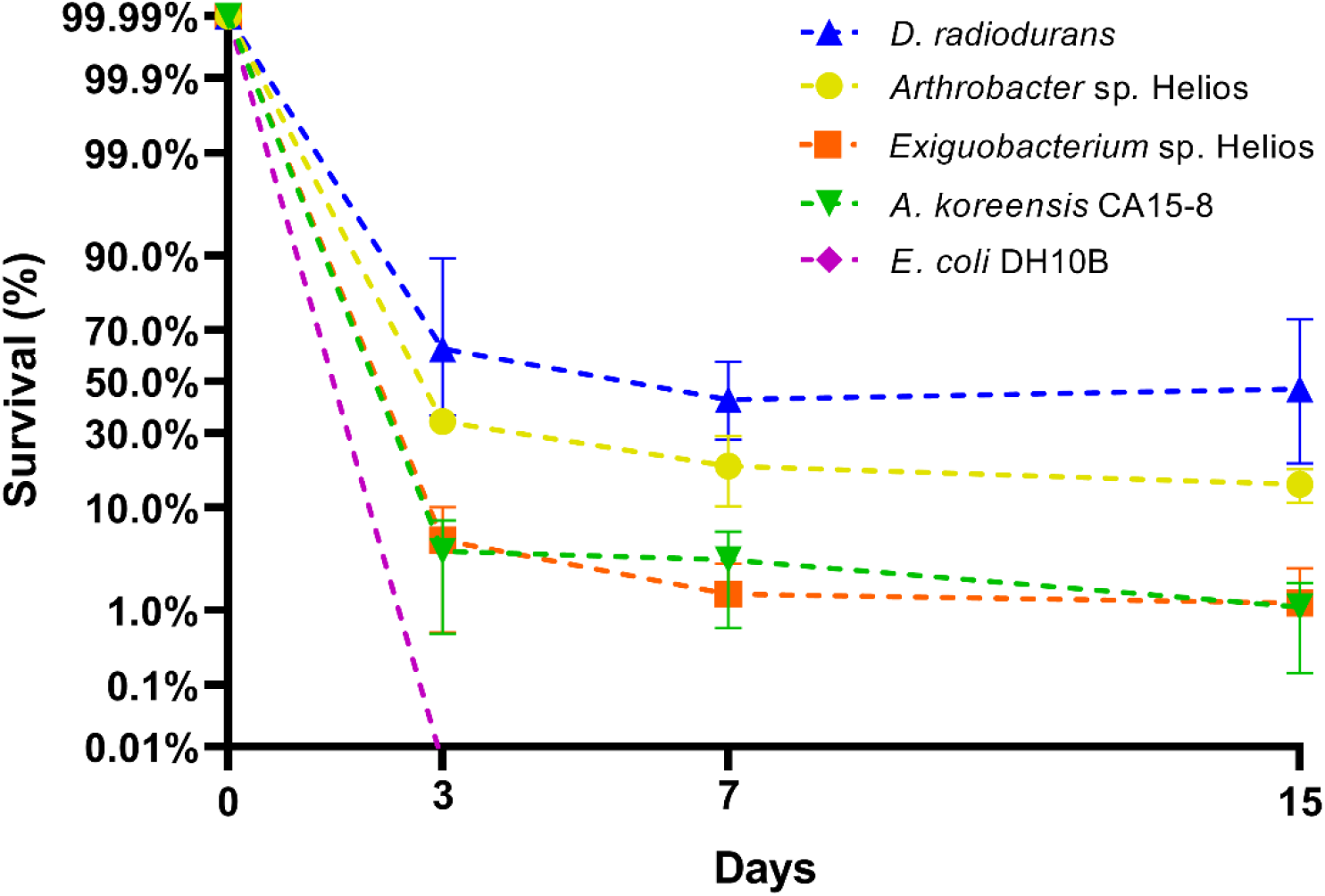
Desiccation tolerance of *Deinococcus radiodurans, Arthrobacter* sp. Helios, *Arthrobacter koreensis* CA15-8 and *Exiguobacterium* sp. Helios cells in exponential phase of growth in LB medium.

The influence of the growth phase on *Arthrobacter* sp. Helios xerotolerance was also checked. A desiccation test was performed with cells from a stationary phase culture and their desiccation tolerance was compared with that of exponential phase cells. Interestingly, *Arthrobacter* sp. Helios showed the same xerotolerance capacity regardless of the growing phase (Fig. 3). In contrast, *Exiguobacterium* sp. Helios and *A. koreensis* CA15-8 survival rate was 10 times lower at the exponential phase than at the stationary phase, suggesting that these strains might use different mechanisms to become adapted to arid environments depending on the growth rates (Fig. 3).

**Figure 3.**
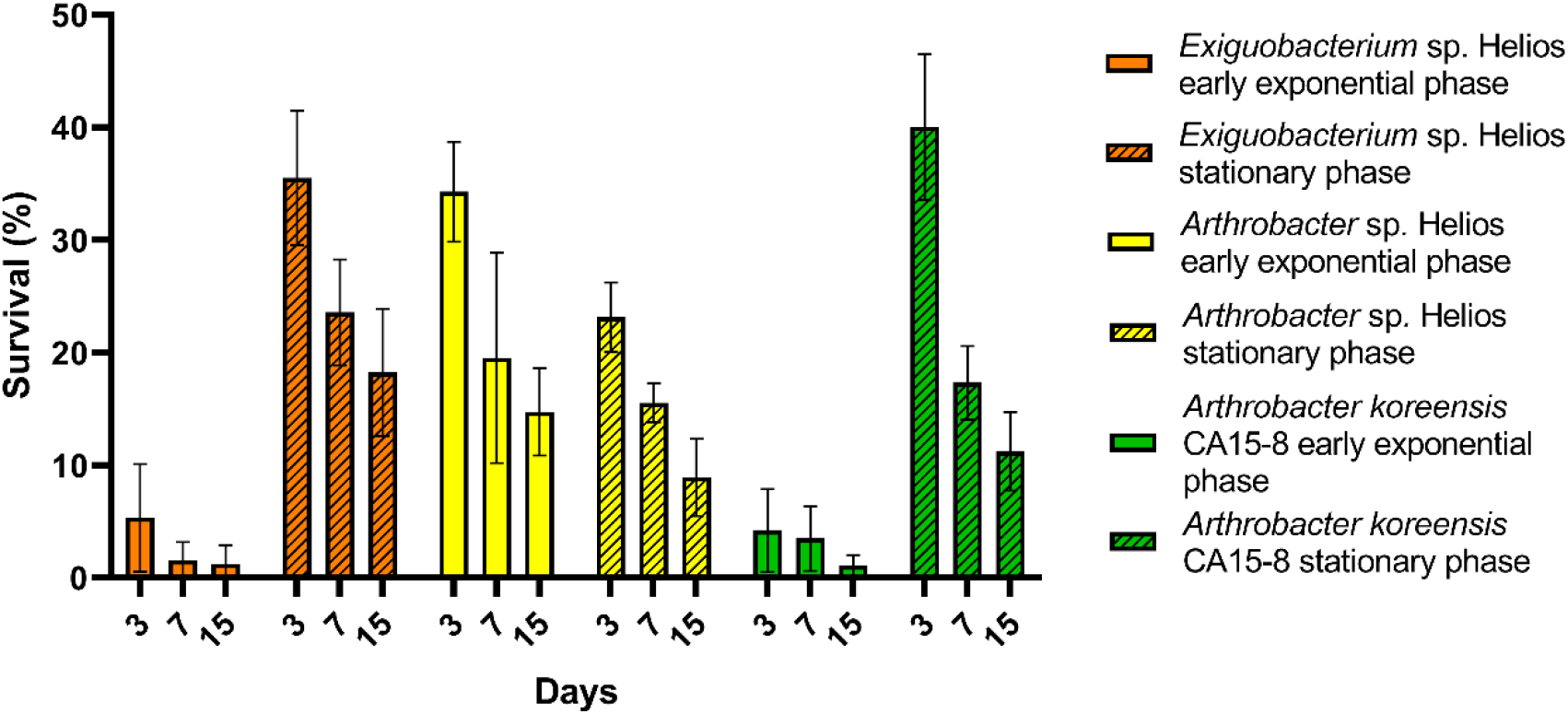
Desiccation tolerance of *Arthrobacter* sp. Helios, *A. koreensis* CA15-8 and *Exiguobacterium* sp. Helios at exponential and stationary growth phases.

### 3.2. Arthrobacter sp. Helios resistance to UV, salinity, metals and metalloids

*Arthrobacter* sp. Helios displays high tolerance to UV radiation. When resistance of *Arthrobacter* sp. Helios to UV radiation was analysed and compared with other strains, it presented a great capacity to survive up to 1500 J/m^2^ of UV radiation (Fig. 4). *Arthrobacter* sp. Helios is also a moderate halotolerant strain since it is able to grow in the presence of 80 g/L NaCl (2-times sea water concentration) (Fig. S1). Furthermore, Table S2 shows that *Arthrobacter* sp. Helios has a moderate resistance to some metals and metalloids. When the strain was grown in LB containing 1 mM selenite at 37 °C, the culture acquired a red colour, suggesting the reduction of selenite to elemental selenium (Fig. S2). No coloration was observed in the absence of bacterial cells, suggesting a role of this strain in selenite reduction. In fact, *Arthrobacter* sp. Helios was able to grow in the presence of selenite up to 150 mM at 37 °C, indicating that the resistance is close to that reported for highly tolerant selenite strains such as *Comamonas testosteroni* S44 (Zheng et al., 2014), *Pseudomonas moraviensis* (Staicu et al., 2015), or *Vibrio natriegens* (Fernández-Llamosas et al., 2017).

**Figure 4.**
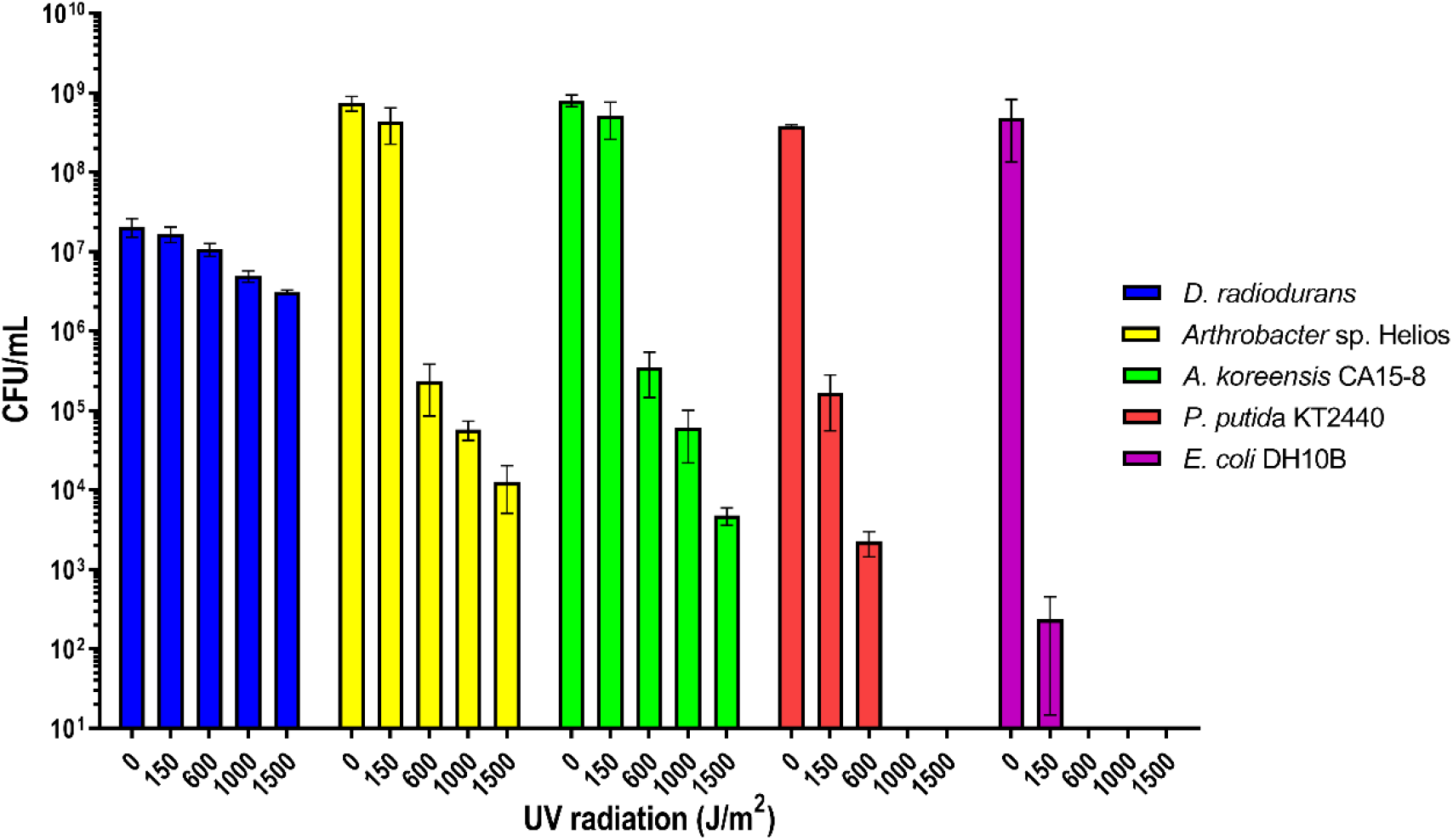
Comparison of the tolerance to UV irradiation of *Arthrobacter* sp. Helios, *A. koreensis* CA15-8, *D. radiodurans, P. putida* KT2440 and *E. coli* DH10B cells in stationary phase grown in LB medium.

Our results suggest that *Arthrobacter* sp. Helios can reduce selenite to elemental Se (0). The red deposits appearance in the growth medium of *Arthrobacter* sp. Helios indicates that, most probably, the selenite is reduced to elemental selenium. Therefore, we checked if this reduction involved the production of selenium nanoparticles, as a proof of concept for developing the Helios strain as a biotechnological tool for the bioproduction of Se nanoparticles (SeNPs). TEM preparations of Helios cultures showed the presence of electron-dense nanospheres in the cells after 24 h of growth at 30 °C in LB containing 1 mM selenite (Figure 5a, 5b). The EDX analysis showed that the nanoparticles presented the specific Se peak (Figure 5c). The selected area electron diffraction (SAED) pattern of the nanoparticles showed a diffuse halo, indicating that selenium is present in its amorphous form (Figure 5c, inset). The shape of SeNPs were spherical with an average size of 229 ± 92 nm (Figure 5d).

**Figure 5.**
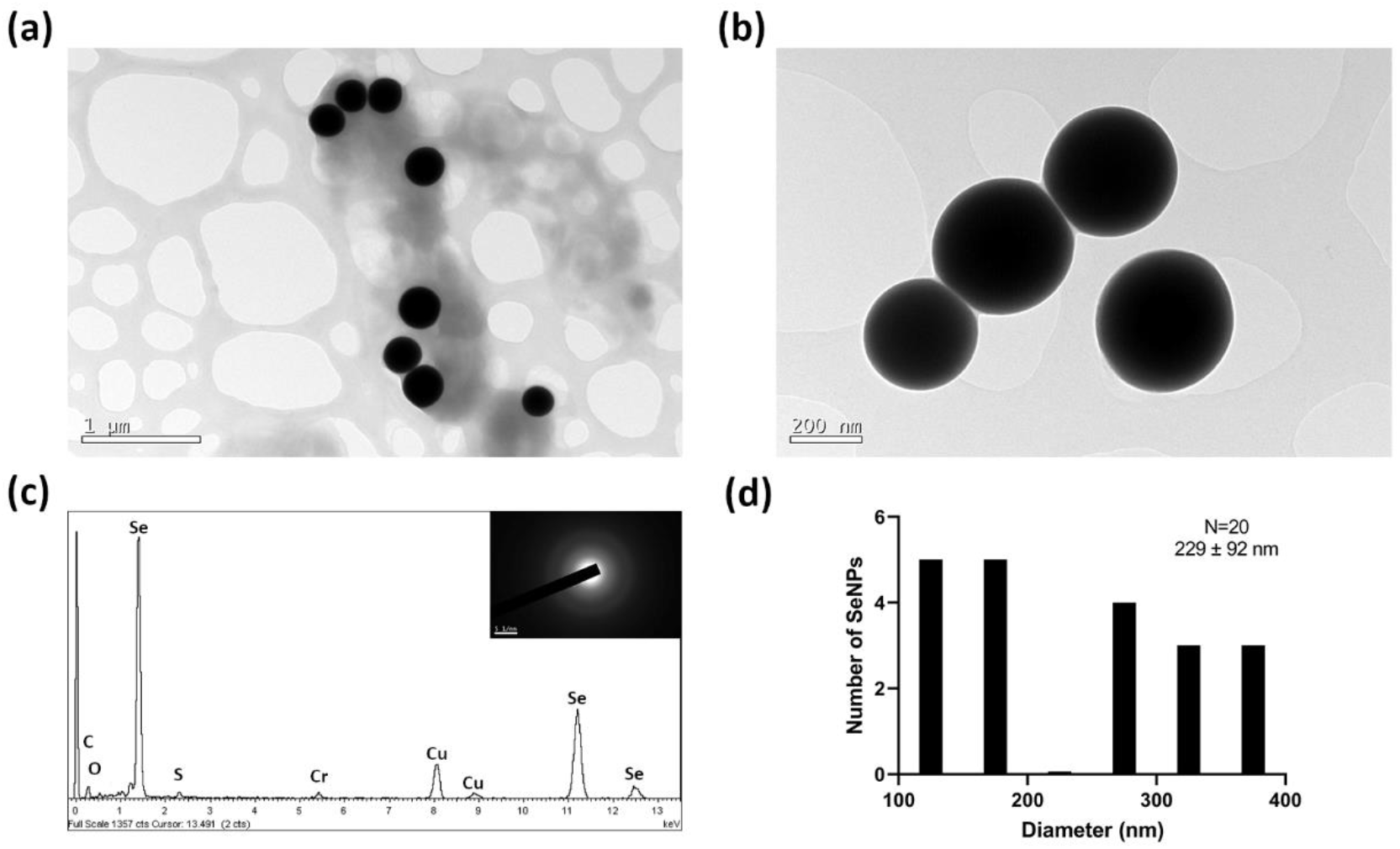
Analysis of the SeNPs produced by *Arthrobacter* sp. Helios. (a, b) TEM images of SeNPs produced by *Arthrobacter* sp. Helios. (c) EDX analysis of one SeNP composition. In the inset is shown SAED pattern of SeNPs. (d) SeNPs size distribution was calculated using the Image J software. Cells were grown for 24 h at 30 °C in LB supplemented with 1 mM sodium selenite.

### 3.3. General features of Arthrobacter sp. Helios genome

The complete genome of *Arthrobacter* sp. Helios consists in a single circular chromosome of 3,895,998 bp, with a 66% GC content and no plasmids. A total of 3,586 genes were predicted of which 2,275 protein-encoding genes were functionally assigned, whereas the remaining genes were predicted as hypothetical proteins. *Arthrobacter* sp. Helios whole genome data was uploaded at the NCBI database under the accession number CP095402.

*Arthrobacter* sp. Helios appears to be ecologically versatile and capable of growing in a variety of carbon sources. The substrate range that *Arthrobacter* sp. Helios can use as a sole carbon source was analysed (Table S3), showing that it can metabolize sugars like glucose, fructose, sucrose, xylose and maltose; organic acids such as acetic acid or pyruvate; aromatic compounds such as protocatechuic acid, 4-hydroxybenzoate, phenyl acetic acid and gentisate, and sterols such as cholesterol. However, Helios strain was not able to catabolize various environmental relevant compounds including pollutants such as phenol, phthalate, terephthalate and isophthalate.

The presence of a large gene cluster, covering a 31 kb region, involved in the synthesis of flagella (*ArtHe_17510-ArtHe_17680*) suggests that this strain is motile (Table S4). The number of alternative RNA polymerase sigma factors is an important strategy of bacteria to successfully face complex environments and induce the response to a particular stress (Lebre et al., 2017). In this sense, *Arthrobacter* sp. Helios contains six σ factors, the same number as *E. coli*, but far of the 62 σ factors found in other actinobacteria like *Streptomyces coelicolor*. The chromosome contains 160 ORFs putatively encoding regulatory proteins belonging to TetR/AcrR (33 proteins), GntR (17 proteins), MarR (13 proteins), LysR (11 proteins), and AraC (5 proteins) family of transcriptional regulators.

Membrane transporters are important for adaptation to low water activity environments to facilitate the uptake of nutrients and the osmolytes required by dehydrated cells. In this sense, *Arthrobacter* sp. Helios genome encodes 349 putative transporters and substrate binding proteins (9.7% of the Helios genome) which is consistent with its environmental versatility.

*Arthrobacter* sp. Helios appears to be well poised to respond to various environmental stresses. The chromosome encodes seven ORFs encoding putative universal-stress proteins (USPs), five ORFs coding cold-shock proteins, and two starvation inducible proteins (Table S4). USP represents a superfamily of proteins whose production is induced in cells in response to several stresses, such as carbon starvation, exposure to UV radiation, osmotic stress, etc. In addition, this bacterium contains one *dnaJ-grpE-dnaK* operon (*ArtHe_12030-12040*) that codes for DnaK and DnaJ chaperons, involved in the response to hyperosmotic and heat shock by preventing the aggregation of stress-denatured proteins. Furthermore, the locus *ArtHe_06610* codes an additional copy of DnaJ chaperone that is not organized in an operon. The genes that code for the essential chaperon GroEL are also present in the genome (*ArtHe_10200* and *ArtHe_00405*). Moreover, *ArtHe_11930* gene codes a putative ClpB chaperone, that is part of a stress-induced multi-chaperone system involved in the recovery of the cell from heat-induced damage, in cooperation with DnaK, DnaJ and GrpE.

*Arthrobacter* sp. Helios is also equipped with many genes encoding ROS scavengers’ enzymes required to survive the internal oxidative stress caused by H_2_O_2_ and other reactive oxygen species that produce extensive damage and cell death. The genome of Helios strain contains genes that code for one superoxide dismutase (*ArtHe_05980*), four catalases (*ArtHe_14485, ArtHe_14120, ArtHe_11075* and *ArtHe_01810*), five peroxidase-coding genes (*ArtHe_12570*, *ArtHe_11330*, *ArtHe_10990*, *ArtHe_08610* and *ArtHe_01040*), two thioredoxin reductases (*ArtHe_14800* and *ArtHe_13320*), and 5 thioredoxin proteins (*ArtHe_16150*, *ArtHe_14805*, *ArtHe_12185*, *ArtHe_11010* and *ArtHe_02295*). Also, there are genes coding for two orthologous of SoxR (*ArtHe_15110* and *ArtHe_01455*), that is known to play an important regulatory role in resistance to oxidative stress.

*Arthrobacter* sp. Helios genome sequence predicts different functions related with the production of osmoprotectants such as trehalose and glycogen, which are known to accumulate under extreme water stress in bacteria protecting the cell against desiccation. The glycogen biosynthetic genes *ArtHe_06185* and *ArtHe_06190* code the glucose-1-phosphate adenyltransferase and the glycogen synthase, respectively. There are up to five different pathways that bacteria may have to accumulate trehalose (Avonce et al., 2006). The presence of several biosynthetic pathways in the same organism suggests a strict requirement to accumulate trehalose under changeable environmental conditions, which could limit substrate availability for each pathway. In this sense, the TPS/TPP pathway involving two enzymatic steps catalysed by trehalose-6-phosphate synthase (TPS) and trehalose-phosphatase (TPP) (Avonce et al., 2006) is present in the Helios strain. *ArtHe_00040* and *ArtHe_00035*, coding TPS and TPP respectively, catalyse the transfer of glucose from UDP-glucose to glucose 6-phosphate forming trehalose 6-phosphate (T6P) and UDP, while TPP dephosphorylates T6P to trehalose and inorganic phosphate. Moreover, *ArtHe_17800* codes a trehalose synthase (TS) which is part of a second biosynthetic pathway in which TS isomerizes the alpha1-alpha4 bond of maltose to an alpha1-alpha1 bond, forming trehalose (Potts 1994; Avonce et al. 2006; Ruhal et al. 2013). A third trehalose biosynthesis pathway is also present in *Arthrobacter* sp. Helios: *ArtHe_10290* and *ArtHe_10285* code two putative maltooligosyl synthase and maltooligosyl trehalohydrolase, respectively, that are involved in the conversion of maltodextrines (maltooligosaccharides, glycogen and starch) to trehalose (Avonce et al., 2006; Ruhal et al., 2013).

Bacteria face osmotic stress by synthesizing or incorporating osmolytes directly from the environment such as glycine betaine (N,N,N-trimethylglycine). One cluster of ABC type glycine/betaine transport genes seems to be present in the Helios strain genome sequence. The *ArtHe_04665-ArtHe_04675* cluster codes two ABC transporter permeases and one ATP binding protein, respectively. Glycine betaine can be synthesized from choline by a two-step pathway with betaine aldehyde as intermediate. The coding genes of a choline uptake protein (*ArtHe_11940*), betaine aldehyde dehydrogenase (*ArtHe_11945*) and choline oxidase (*ArtHe_11950*) are forming a putative operon in the Helios strain with an additional gene coding a choline uptake protein (*ArtHe_13245*). Finally, Helios strain contains two putative aquaporin Z coding genes (*ArtHe_08770 and ArtHe_06700*), that are known to modulate water fluxes and therefore could have a main role in cell homeostasis.

The operon *ArtHe_11495-11510* codes the three subunits of the nitrate respiratory reductase and the cofactor assembly chaperone. The presence of this enzyme suggests that *Arthrobacter* sp. Helios has the ability to use nitrate (NO_3_) during anaerobic respiration and reduce it to nitrite (NO_2_).

To explore possible biotechnological applications of *Arthrobacter* sp. Helios, we searched possible genes that could be involved in plant growth promoting. In this sense, *ArtHe_14430* codes a putative 1-aminocyclopropane-1-carboxylate (ACC) deaminase that is involved in lowering plant ethylene levels, often a result of various stresses. The ACC deaminase synergistically interacts with the plant and bacterial auxin indole-3-acetic acid (IAA). *ArtHe_01245* gene, annotated as amidase, codes a protein very similar to the indolacetamide hydrolase involved in IAA biosynthesis in *Agrobacterium tumefaciens* and *Rhodococcus* sp. N774 (Hashimoto et al., 1991). Nitrogen fixation is one of the most remarkable plant-growth-promoting properties among plant colonising bacteria, and this activity is coded in the *nif* genes (Mus et al., 2018). However, no *nif*-homologous genes were found in the *Arthrobacter* sp. Helios genome, suggesting that this strain is not able to fix atmospheric nitrogen into ammonia.

### 3.4. Identification of differentially expressed genes in the presence of PEG6000

To analyse the desiccation resistance mechanisms of *Arthrobacter* sp. Helios under low external water activity conditions, we used PEG6000 since these molecules are too large to pass the cytoplasmic membrane and can simulate effectively a matric stress condition to bacteria without causing the toxic effects due to high concentrations of ions used to recreate low water activity conditions (Holden, 2001). First, to set up the culture conditions to perform transcriptomic analyses, *Arthrobacter* sp. Helios was cultured in the presence of increasing concentrations of PEG6000. Figure 6a shows that bacterial growth is not impaired at PEG concentrations up to 20% (w/v). However, at 30% and 35% of PEG the duplication rates were lower, i.e., 11 and 41 h, respectively, with a decrease on the biomass at the end of the growth curve (Fig. 6b). No growth in the presence of 40% PEG6000 or higher concentrations was observed. Based on the results, the conditions selected to find differences in the transcriptome were 0% (PEG0), 10% (PEG10) and 35% PEG6000 (PEG35).

**Figure 6.**
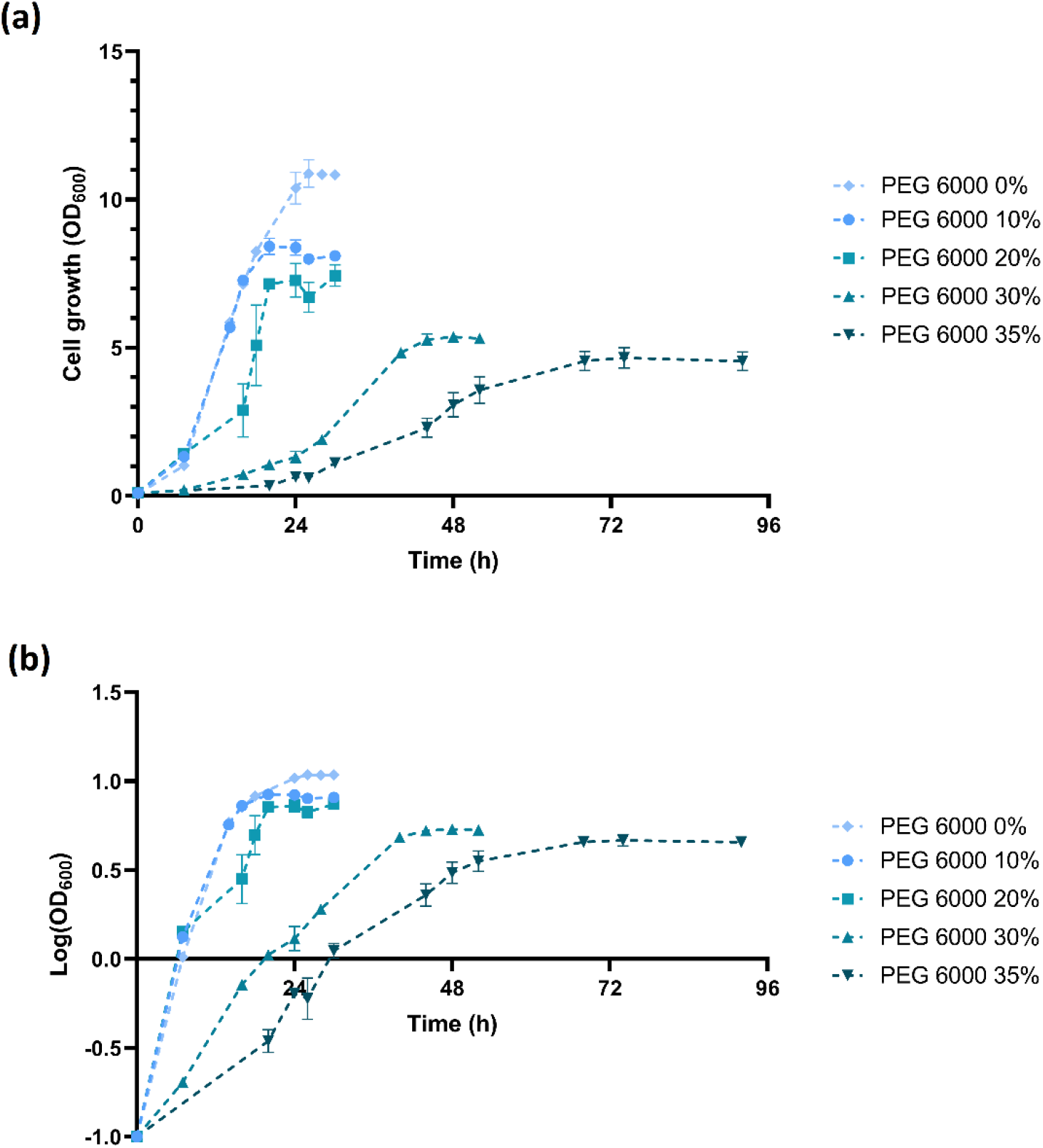
Growth of *Arthorbacter* sp. Helios in LB medium with increasing concentrations of PEG 6000. (a) Growth curve (b) Semilogarithmic representation. Mean and standard deviation of three replicas are represented.

*Arthrobacter* sp. Helios cells were collected in the middle of the exponential growth phase in LB medium with 0%, 10% and 35% PEG6000 and RNA was extracted to perform the *de novo* transcriptome analysis. A total of 324 differentially expressed genes (DEG) were identified in the PEG35 vs. PEG0 (control) conditions. Among them, 184 genes were upregulated and 140 were downregulated. The distribution of DEG in both conditions according to their log_2_FC and - log_10_FDR are represented in the volcano maps shown in Figure 7. Meanwhile, 105 DEG were identified in the PEG10 condition compared with the control condition PEG0, from which 52 were upregulated and 53 were downregulated. The comparison of three conditions (PEG0 vs PEG10 and PEG35) shared 29 upregulated and 13 downregulated genes (Fig. 8). The principal component analysis (Fig. S3) showed a high correspondence between the three biological replicates of each condition, and each group was markedly separated from the others. These results proved that not only the RNA-seq data were highly reproducible but also that there was a unique gene expression at different levels of drought stress.

**Figure 7.**
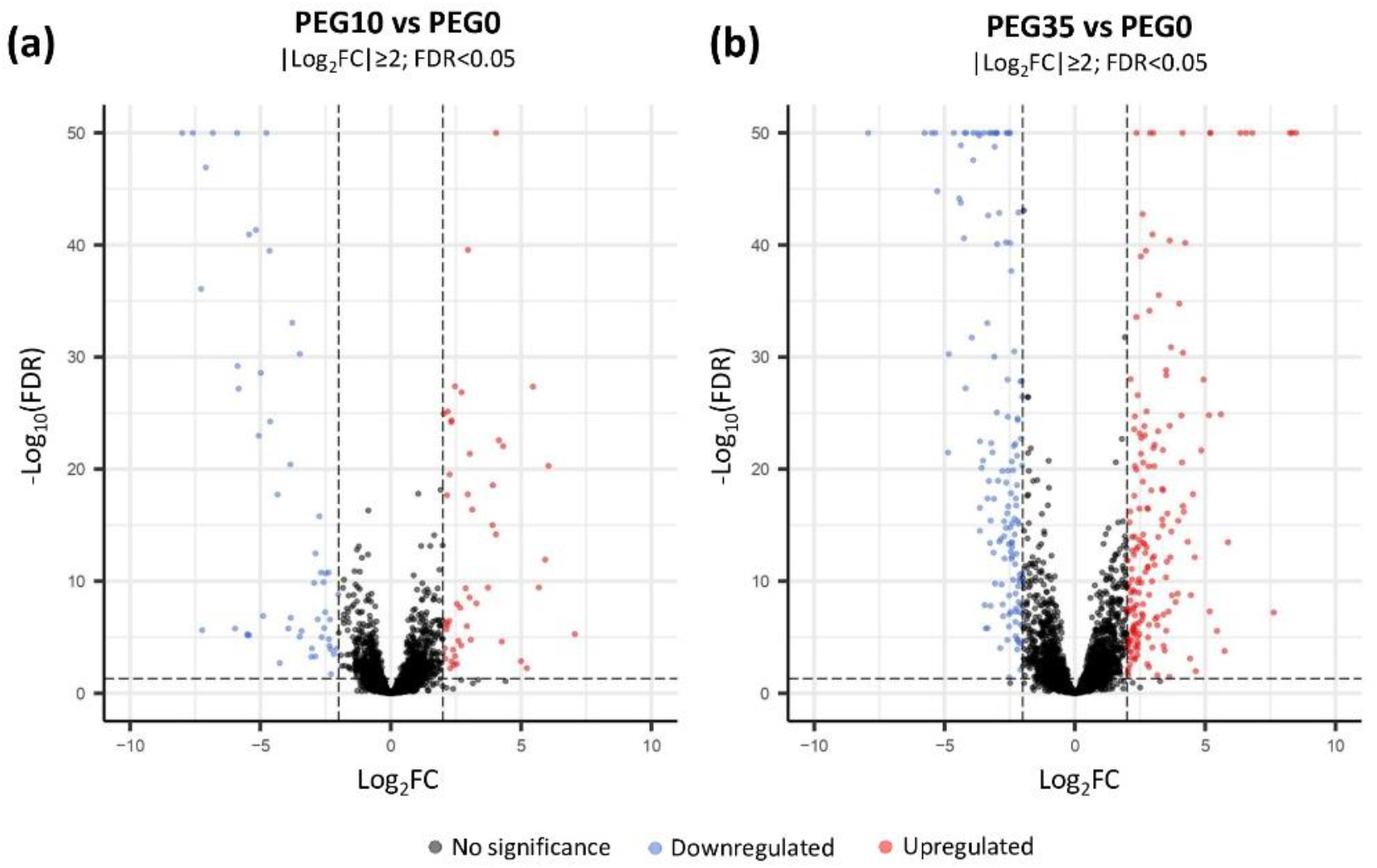
Volcano maps showing the distribution of DEGs according to their log_2_FC and -log_10_FDR. (a) DEGs found in PEG10 condition compared to control condition PEG0. (b) DEGs found in PEG35 condition versus the control condition PEG0. Grey spots represent genes with a non-significant expression change, while blue and red circles represent down regulated and up regulated genes, respectively. Genes are considered differentially expressed when |log_2_FC| ≥2 and FDR<0.05.

**Figure 8.**
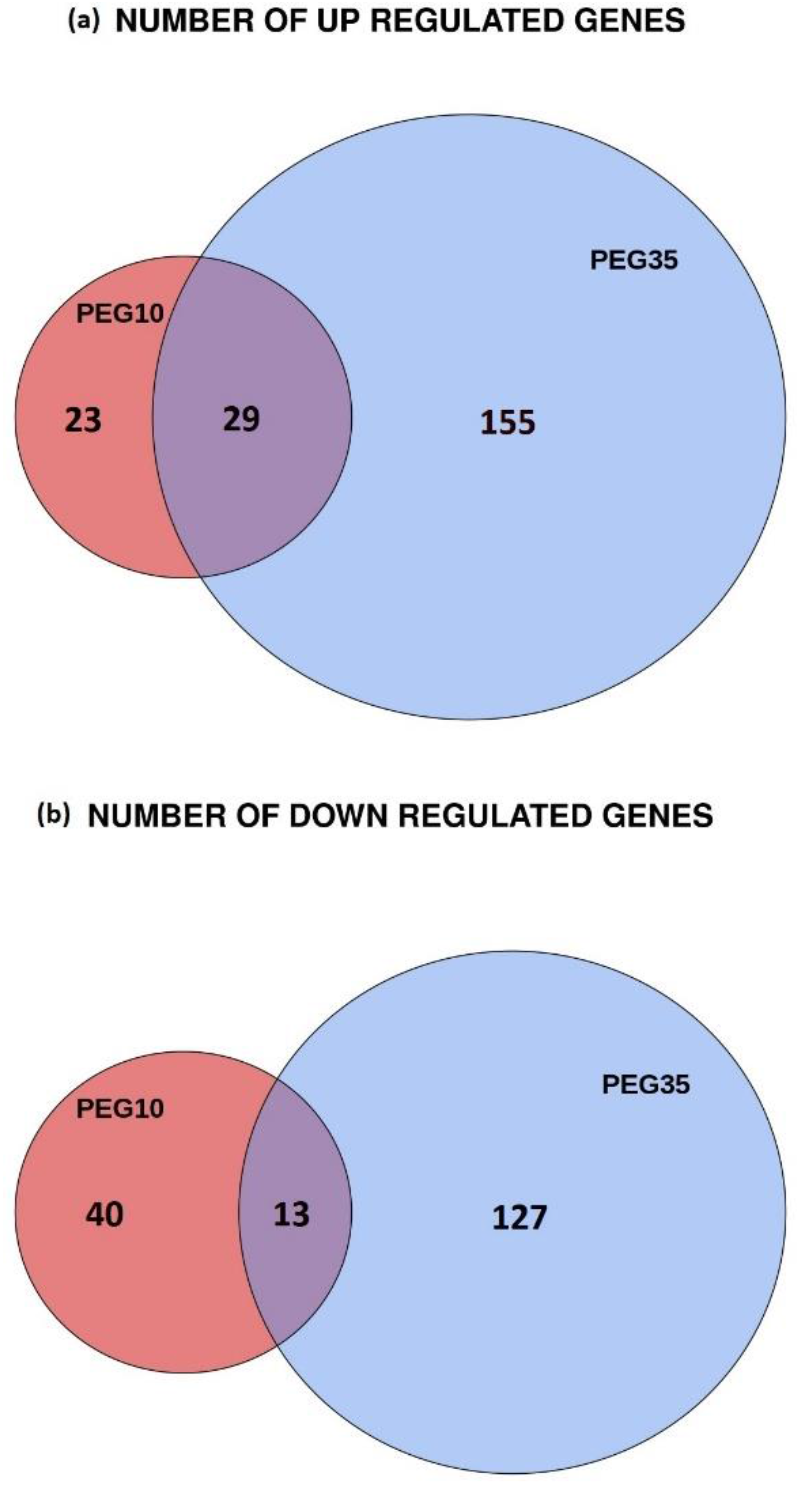
Venn diagrams summarizing the DEGs of *Arthrobacter* sp. Helios in the simulated drought versus control conditions. (a) Number of upregulated genes in PEG10 vs. PEG0 (red circle) and PEG35 vs. PEG0 (blue circle) comparisons. (b) Number of downregulated genes in PEG10 vs. PEG0 (red circle) and PEG35 vs. PEG0 (blue circle) conditions. Genes are considered differentially expressed when |log_2_FC| ≥2 and FDR<0.05.

COG and KEGG databases were used to functionally annotate DEGs. Although DEGs were assigned to all COG categories, some of them contained a significantly higher proportion of upregulated or downregulated genes (Fig. 9). Using a hypergeometric test in the condition PEG35 vs PEG0, the categories having statistically enriched upregulated genes were energy production and conversion (C), amino acid transport and metabolism (E) and inorganic ion transport and metabolism (P). In contrast, only cell motility (N) category was enriched with downregulated genes (Fig. 9a). Many of the changes observed in PEG35 were already detected in PEG10 such as those assigned to the inorganic ion transport and metabolism category (P). However, we found the genes assigned to category C and secondary metabolites biosynthesis, transport and catabolism category (Q) to be downregulated (Fig. 9b). These findings suggested that the ion transport and metabolism seemed to be a common bacterial response against low and high drought stress, although there are some differences depending on the level of stress (PEG10 vs PEG35) (Table S5). According to KEGG annotations (Table S6), ABC transporter genes in both conditions studied presented a significant upregulation, whereas flagella assembly and nitrogen metabolism associated genes showed a significant downregulation.

**Figure 9.**
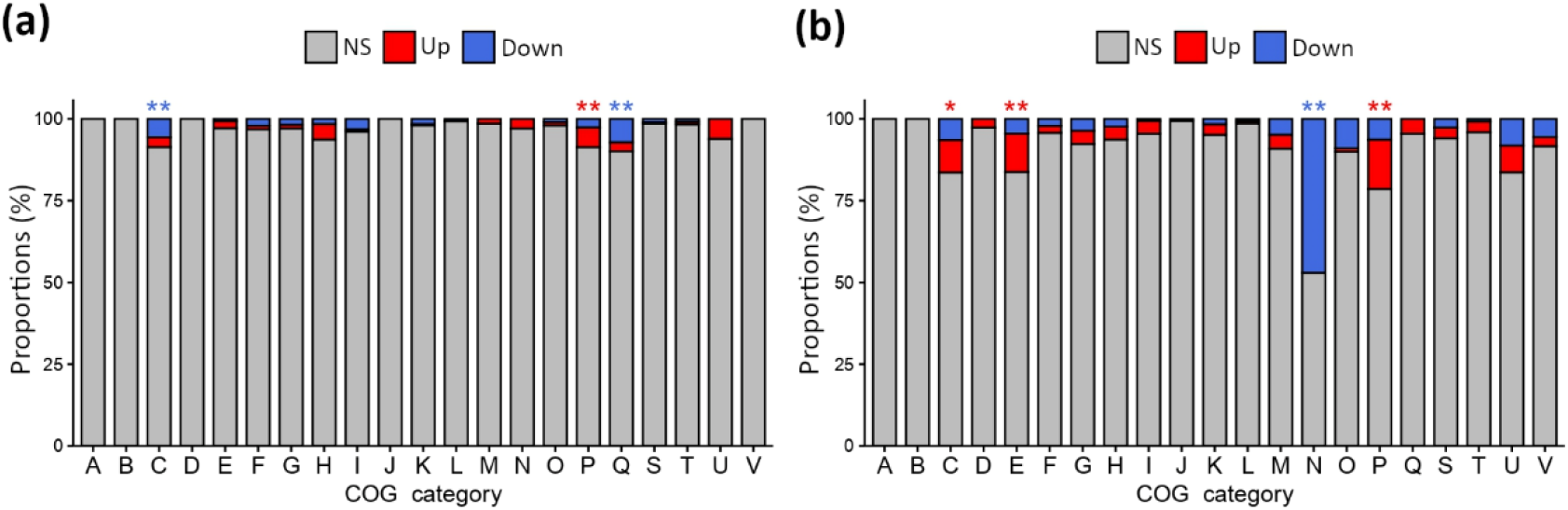
COG distribution of *Arthrobacter* sp. Helios DEGs found under simulated desiccation and control conditions. (a) Proportion of DEGs in each COG category when PEG10 condition was compared against control condition PEG0. (b) Proportion of DEGs in each COG category when PEG35 condition was compared against PEG0 condition. The proportions of non-significant (grey), upregulated (red) and downregulated (blue) genes are shown. A: RNA processing and modification; B: Chromatin structure and dynamics; C: Energy production and conversion; D: Cell cycle control, cell division, chromosome partitioning; E: Amino acid transport and metabolism; F: Nucleotide transport and metabolism; G: Carbohydrate transport and metabolism; H: Coenzyme transport and metabolism; I: Lipid transport and metabolism; J: Translation, ribosomal structure and metabolism; K: Transcription; L: Replication, recombination and repair; M: Cell wall/membrane/envelop biogenesis; N: Cell motility; O: Post-translational modification, protein turnover, chaperones; P: Inorganic ion transport and metabolism; Q: Secondary metabolites biosynthesis, transport and catabolism; R: General function prediction only; S: Function unknown; T: Signal transduction mechanisms; U: Intracellular trafficking, secretion and vesicular transport; V: Defence mechanisms. A hypergeometric test was performed to check whether genes were significantly overrepresented in each category (*=p.adjust<0.05; **=p.adjust<0.01).

Bacteria have developed different strategies to overcome osmotic stress and retain water inside the cell when the environment turns hypertonic. As shown in Table 1, *Arthrobacter* sp. Helios facing water shortage induced the expression of *ArtHe_04675* and *ArtHe_04665*, genes coding glycine betaine ABC transporters involved in the uptake of this compatible solute. The highest induction (>50 fold) was observed in the *kdpA* (*ArtHe_17390*), *kdpB* (*ArtHe_17385*) and *kdpF* (*ArtHe_17395*) genes of the *kdpFABC* operon encoding a potassium transport ATPase, which catalyses the hydrolysis of ATP coupled with the electrogenic transport of potassium into the cytoplasm. The expression of the *ArtHe_12610* gene coding a CapA family protein, similar to *Bacillus anthracis* capsule polysaccharide biosynthesis, is induced 8-fold. The presence of PEG also induced 5-fold the expression of *ArtHe_12305* gen, that codes a mechanosensitive ion channel family protein that participates in cellular homeostasis. As expected, the aquaporin coding genes *ArtHe_06700* and *ArtHe_08770*) are downregulated (2 and 5-fold, respectively) most probably to avoid the loss of water by a facilitated diffusion through this channel. Remarkably, no other osmolyte biosynthesis genes, such as those involved in trehalose production, were significantly over-expressed in these conditions (Table S7).

**Table 1.**
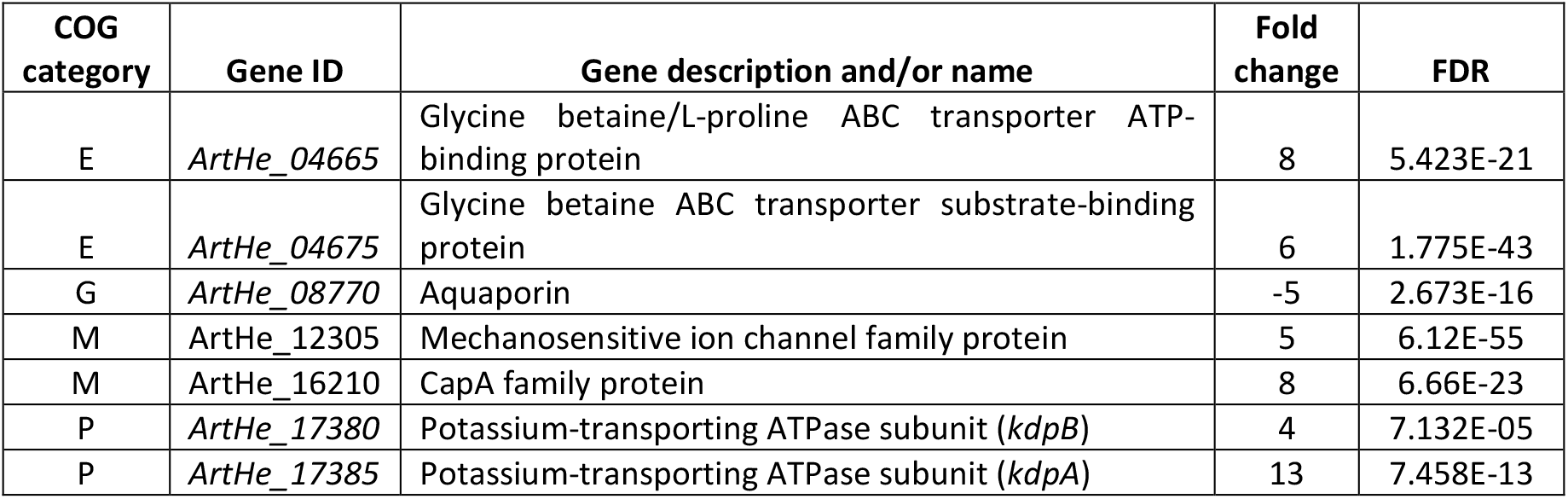
DEGs related to osmotic stress in PEG35 vs. PEG0

| COG category | Gene ID | Gene description and/or name | Fold change | FDR |
| --- | --- | --- | --- | --- |
| E | *ArtHe_04665* | Glycine betaine/L-proline ABC transporter ATP-binding protein | 8 | 5.423E-21 |
| E | *ArtHe_04675* | Glycine betaine ABC transporter substrate-binding protein | 6 | 1.775E-43 |
| G | *ArtHe_08770* | Aquaporin | -5 | 2.673E-16 |
| M | *ArtHe_12305* | Mechanosensitive ion channel family protein | 5 | 6.12E-55 |
| M | *ArtHe_16210* | CapA family protein | 8 | 6.66E-23 |
| P | *ArtHe_17380* | Potassium-transporting ATPase subunit (*kdpB*) | 4 | 7.132E-05 |
| P | *ArtHe_17385* | Potassium-transporting ATPase subunit (*kdpA*) | 13 | 7.458E-13 |

Along with inorganic ions transporters, amino acid and peptide ABC transporters were also massively induced in the Helios strain (Table 2). Most of them are grouped in gene clusters in *Arthrobacter* sp. Helios genome and their induction was paired with the upregulation of an aminotransferase, a serine ammonia-lyase, *hyuA* and *dapA*. The *hyuA* gene encodes a hydantoin racemase, involved in the racemization of amino acid precursors, and *dapA* encodes a dihydrodipicolinate synthase, a key enzyme in lysine biosynthesis via diaminopimelic acid.

**Table 2.** DEGs involved in amino acid and nitrogen metabolism in PEG35 vs. PEG0

| COG category | Gene ID | Gene description and/or name | Fold change | FDR |
| --- | --- | --- | --- | --- |
| E | *ArtHe_00090* | ABC transporter substrate-binding protein | 37 | 8.9295E-70 |
| E | *ArtHe_00190* | ABC transporter ATP-binding protein | 5 | 2.6902E-34 |
| E | *ArtHe_01080* | ABC transporter ATP-binding protein | 7 | 0.00247634 |
| E | *ArtHe_01125* | Aminotransferase class III-fold pyridoxal phosphate-dependent enzyme | 6 | 5.0422E-05 |
| E | *ArtHe_01270* | ABC transporter permease | 5 | 5.0422E-05 |
| K/O | *ArtHe_01770* | Urease accessory protein (*ureG*) | -9 | 9.3437E-31 |
| O | *ArtHe_01775* | Urease accessory protein (*ureF*) | -13 | 3.293E-15 |
| E | *ArtHe_01780* | Urease subunit alpha | -8 | 8.9318E-26 |
| E | *ArtHe_01790* | Urease subunit gamma | -5 | 3.0468E-12 |
| E | *ArtHe_04130* | Urease subunit gamma | -5 | 2.094E-21 |
| E | *ArtHe_04140* | Urease subunit beta | -7 | 9.8499E-15 |
| E | *ArtHe_04145* | Urease subunit alpha | -9 | 4.8753E-23 |
| O | *ArtHe_04150* | Urease accessory protein | -6 | 1.3713E-20 |
| K/O | *ArtHe_04155* | Urease accessory protein (*ureG*) | -5 | 1.7988E-17 |
| O | *ArtHe_04160* | Urease accessory protein (*ureD*) | -4 | 1.9348E-23 |
| E | *ArtHe_04170* | Urea transporter | -5 | 4.3705E-07 |
| E | *ArtHe_04615* | Hydantoin racemase (*hyuA*) | 5 | 1.1471E-14 |
| K | *ArtHe_08230* | P-II family nitrogen regulator | -29 | 3.3462E-22 |
| P | *ArtHe_08235* | Ammonium transporter | -19 | 2.5332E-41 |
| E | *ArtHe_11200* | APC family permease | 7 | 7.4576E-35 |
| P | *ArtHe_11490* | NarK/NasA family nitrate transporter | -5 | 2.0897E-38 |
| C | *ArtHe_11495* | Respiratory nitrate reductase subunit gamma | -10 | 4.0125E-14 |
| C | *ArtHe_11500* | Nitrate reductase molybdenum cofactor assembly chaperone | -5 | 7.0658E-16 |
| C | *ArtHe_11505* | Nitrate reductase subunit beta | -5 | 3.1799E-31 |
| C | *ArtHe_11510* | Nitrate reductase subunit alpha | -4 | 4.8241E-21 |
| S | *ArtHe_12365* | M81 family metallopeptidase | 59 | 3.4091E-14 |
| E | *ArtHe_12370* | D-serine ammonia-lyase | 8 | 2.4774E-07 |
| E | *ArtHe_12530* | ABC transporter ATP-binding protein | 5 | 5.4086E-21 |
| E | *ArtHe_13820* | ATP-binding cassette domain-containing protein | 13 | 3.7509E-15 |
| E | *ArtHe_13825* | ABC transporter ATP-binding protein | 16 | 4.1653E-16 |
| E | *ArtHe_13830* | Branched-chain amino acid ABC transporter permease | 18 | 6.5006E-17 |
| E | *ArtHe_13835* | Branched-chain amino acid ABC transporter permease | 13 | 1.346E-24 |
| E | *ArtHe_13840* | ABC transporter substrate-binding protein | 18 | 4.1367E-31 |
| E | *ArtHe_14305* | GMC family oxidoreductase | 95 | 1.14E-173 |
| C | *ArtHe_14310* | Aldehyde dehydrogenase family protein | 112 | 4.8281E-75 |
| E | *ArtHe_14315* | APC family permease | 305 | 4.265E-174 |
| K | *ArtHe_14320* | Helix-turn-helix domain-containing protein | 81 | 1.292E-194 |
| Q | *ArtHe_14340* | Primary-amine oxidase | 323 | 0 |
|  | *ArtHe_14345* | Hypothetical protein | 359 | 3.8362E-58 |
| E | *ArtHe_16215* | Amino acid ABC transporter ATP-binding protein | 10 | 1.0563E-15 |
| E | *ArtHe_16220* | Amino acid ABC transporter permease | 5 | 1.0953E-07 |
| E | *ArtHe_16565* | Amino acid ABC transporter ATP-binding protein | 4 | 0.00272718 |
| E | *ArtHe_16635* | Dihydrodipicolinate synthase family protein (*dapA*) | 10 | 5.8743E-19 |

*ArtHe_12365* coding a putative metallopeptidase is one of the most induced genes (Table 2). It has been hypothesized that the increase of the proteolytic activity could play an essential role removing damaged proteins and release amino acids that would provide osmolytes to compensate the osmotic shock. (Frees et al., 2007; Ghedira et al., 2018).

The greatest induction (up to 321-fold) in PEG35 was observed in a 11 kb gene cluster spanning from *ArtHe_14305* to *ArtHe_14345* genes that is putatively involved in the transport and catabolism of phenylethylamine (PEA) (Table 2). PEA is a biogenic amine that is catabolized by some bacteria (Luengo and Olivera, 2020). In general, it is described that biogenic amines are involved in chromosomal and ribosomal organization, DNA replication/translation and in the regulation of RNA and protein synthesis in prokaryotes (Perry and Fetherston, 2007; Luengo and Olivera, 2020). Polyamines carry a net positive charge at physiological pH that forms electrostatic bonds with negatively charged macromolecules such as DNA/RNA, for maintaining a stable conformation. Generally, polyamines have been broadly implicated in cell growth due to their ability to interact with nucleic acids and protein translation machinery (Tabor and Tabor, 1985; Dever and Ivanov, 2018). They have a protective role against radiations and oxidative stress and participate in biofilm formation. These molecules can act as free radical scavengers by interaction and subsequent inactivation with these molecules. (Perry and Fetherston, 2007). Moreover, spermidine/putrescine ABC transporters genes were found upregulated in *Arthrobacter* sp. Helios (Table 3). Putrescine and spermidine are polyamines that are essential components for deoxyribonucleic acid (DNA) packaging during the cell cycle.

**Table 3.** DEGs involved in DNA protection and general stress in PEG35 vs. PEG0

| COG category | Gene ID | Gene description and/or name | Fold change | FDR |
| --- | --- | --- | --- | --- |
| L/U | *ArtHe_08110* | DNA-processing protein DprA | 5 | 0.00039694 |
| T | *ArtHe_11205* | Universal stress protein | 8 | 4.439E-89 |
| E | *ArtHe_12855* | ABC transporter ATP-binding protein | 35 | 0.00015476 |
| E | *ArtHe_12860* | Extracellular solute-binding protein | 15 | 5.9055E-05 |
| E | *ArtHe_12865* | ABC transporter permease | 11 | 1.2996E-09 |
| E | *ArtHe_12870* | ABC transporter permease | 11 | 1.5486E-25 |
| F | *ArtHe_13950* | 8-oxoguanine deaminase | 4 | 9.251E-08 |
| T | *ArtHe_16365* | Universal stress protein | 3 | 0.01644528 |

The 8-oxoguanine deaminase coding gene was found upregulated in *Arthrobacter* sp. Helios (Table 3). The 8-oxoguanine deaminase is known to play a critically important role in the DNA repair activity for oxidative damage and catalyses the conversion of 8-oxoguanine, formed by the oxidation of guanine residues within DNA by reactive oxygen species, in urate and ammonia (De Rosa et al. 2021).

Only the *ArtHe_11205* and *ArtHe_16365* genes encoding two of the six universal stress proteins (USPs) annotated in the Helios genome are induced 8- and 3-fold, respectively (Table 3). None of the genes coding cold shock proteins are being induced in the presence of PEG. In contrast, the genes coding GroEL (*ArtHe_00405* and *ArtHe_10200*) and ClpB (*ArtHe_11930*) chaperonins are slightly induced (2-fold) (Table S7). *DprA* gene (*ArtHe_08110*), encoding a DNA-processing protein that binds ssDNA and loads RecA during transformation, was also greatly induced, suggesting some kind of DNA protection role.

Iron scavenging is known to be crucial for bacterial survival. One mechanism for iron scavenging is the siderophore-mediated acquisition through specific receptor and transport systems and siderophores have been extensively reported to reduce oxidative stress in microorganisms (Khan et al., 2018)*. Arthrobacter* sp. Helios transcriptomic data shows how several iron and siderophore ABC transporter genes are highly upregulated in drought conditions. The different clusters encoding the iron transporter components have a variable induction ranging from 4-fold to 197-fold (Table 4). *ArtHe_12765* gene that codes a heme-oxygenase, probably involved in iron reutilization, is induced by 13-fold, while *ArtHe_10780* coding the iron storage protein ferritin is downregulated 7-fold.

**Table 4.** DEGs related to iron homeostasis in PEG35 vs. PEG0

| COG category | Gene ID | Gene description and/or name | Fold change | FDR |
| --- | --- | --- | --- | --- |
| P | *ArtHe_05605* | Iron chelate uptake ABC transporter family permease subunit | 21 | 8.2729E-05 |
| P | *ArtHe_05610* | ABC transporter substrate-binding protein | 9 | 7.2506E-09 |
| P | *ArtHe_10210* | Siderophore ABC transporter substrate-binding protein | 49 | 6.2336E-08 |
| P | *ArtHe_10215* | ABC transporter permease | 6 | 2.3039E-06 |
| P | *ArtHe_10220* | Iron chelate uptake ABC transporter family permease subunit | 5 | 4.7516E-06 |
| P | *ArtHe_10225* | ATP-binding cassette domain-containing protein | 5 | 6.94E-15 |
| P | *ArtHe_10780* | Ferritin | -7 | 1.3086E-25 |
| H/P | *ArtHe_12265* | ABC transporter ATP-binding protein | 6 | 1.5863E-07 |
| P | *ArtHe_12270* | Iron chelate uptake ABC transporter family permease subunit | 7 | 0.00081522 |
| P | *ArtHe_12310* | Iron-siderophore ABC transporter substrate-binding protein | 5 | 1.9942E-10 |
| P | *ArtHe_12765* | Biliverdin-producing heme oxygenase | 13 | 5.0593E-05 |
| P | *ArtHe_13015* | Fe<sup>2+</sup>-enterobactin ABC transporter substrate-binding protein | 6 | 2.659E-06 |
| P | *ArtHe_13020* | Iron ABC transporter permease | 5 | 5.1296E-08 |
| U | *ArtHe_13025* | Iron chelate uptake ABC transporter family permease subunit | 5 | 0.00074103 |
| H/P | *ArtHe_13030* | ATP-binding cassette domain-containing protein | 12 | 7.1902E-05 |
| P | *ArtHe_13035* | Siderophore-interacting protein | 11 | 1.1931E-07 |
| P | *ArtHe_13040* | DUF3327 domain-containing protein | 4 | 1.8947E-09 |
| H/P | *ArtHe_13545* | ABC transporter ATP-binding protein | 198 | 2.8605E-07 |
| P | *ArtHe_13550* | Iron-siderophore ABC transporter substrate-binding protein | 15 | 1.6635E-09 |
| P | *ArtHe_13555* | Iron ABC transporter permease | 8 | 1.1208E-20 |
| P | *ArtHe_16990* | ABC transporter substrate-binding protein | 36 | 8.6443E-07 |
| P | *ArtHe_16995* | Iron ABC transporter permease | 9 | 5.10E-08 |

Although the genome of *Arthrobacter* sp. Helios is fully equipped with many genes coding ROS scavengers’ enzymes, only one peroxidase-coding gene *ArtHe_10990* is induced 3-fold in PEG condition. Five genes coding thioredoxins (*ArtHe_01455, ArtHe_02295, ArtHe_11010 ArtHe_12185, ArtHe_14805*) did not change in PEG conditions (Table S7).

*Arthrobacter* sp. Helios under simulated drought stress showed almost the whole flagellum biosynthesis cluster downregulated, as shown in Table 5. The genes *fliD, fliF, fliG, fliM, fliN, fliR, fliS, flgB* and *flgC* showed a diminished transcription ranging from 5 to 12-fold, being the flagellin encoding gene the most downregulated one with a 21-fold change.

**Table 5.** DEGs involved in flagellum biosynthesis in PEG35 vs. PEG0

| COG category | Gene ID | Gene description and/or name | Fold change | FDR |
| --- | --- | --- | --- | --- |
| N | *ArtHe_17535* | Flagellin | -21 | 1.7263E-44 |
| N | *ArtHe_17540* | Flagellar filament capping protein (*fliD*) | -13 | 1.7689E-50 |
| N/O/U | *ArtHe_17545* | Flagellar export chaperone (*fliS*) | -9 | 3.9897E-16 |
| N | *ArtHe_17560* | Flagellar biosynthesis protein (*flgB*) | -11 | 1.6555E-06 |
| N | *ArtHe_17565* | Flagellar basal-body rod protein (*flgC*) | -6 | 9.0718E-08 |
| N | *ArtHe_17570* | Flagellar hook-basal body complex protein (*fliE*) | -11 | 1.3726E-08 |
| N/U | *ArtHe_17575* | Flagellar M-ring protein (*fliF*) | -7 | 1.3841E-43 |
| N | *ArtHe_17580* | Flagellar motor switch protein (*fliG*) | -5 | 8.9054E-23 |
| N | *ArtHe_17620* | Flagellar FlbD family protein | -8 | 1.6667E-10 |
| N | *ArtHe_17625* | MotA/TolQ/ExbB proton channel family protein | -4 | 7.9446E-07 |
| N | *ArtHe_17630* | Flagellar motor protein (*motB*) | -4 | 2.0114E-11 |
| N | *ArtHe_17635* | Flagellar motor switch protein (*fliM*) | -5 | 8.4607E-14 |
| N/U | *ArtHe_17640* | Flagellar motor switch protein (*fliN*) | -5 | 1.0013E-12 |
| N | *ArtHe_17645* | FliO/MopB family protein | -6 | 2.0956E-17 |
| N | *ArtHe_17650* | Flagellar type III secretion system pore protein (*fliP*) | -9 | 3.2463E-22 |
| N/U | *ArtHe_17660* | Flagellar biosynthetic protein (*fliR*) | -5 | 7.4035E-11 |

*Arthrobacter* sp. Helios grown in PEG10, i.e., under moderate drought stress, showed a lower number of DEGs than in PEG35, 52 DEGs were upregulated and 53 down regulated (Fig. 8). However, some of the 29 upregulated DEGs in common with the PEG35 condition showed a higher fold change in PEG10 (Table 6). Among those genes, we found potassium transporters, spermidine/putrescine transporters and a CapA family protein. Besides, this drought condition significantly induced the transcription of genes involved in riboflavin biosynthesis, a response not seen at all in PEG35.

**Table 6.** DEGs in PEG10 vs PEG0

| COG category | Gene ID | Gene description and/or name | Fold change | FDR |
| --- | --- | --- | --- | --- |
| H | *ArtHe_05120* | Riboflavin synthase (*ribC*) | 5 | 0.00013257 |
| H | *ArtHe_05125* | 3,4-dihydroxy-2-butanone-4-phosphate synthase (*ribB*) | 6 | 2.1007E-05 |
| H | *ArtHe_05130* | GTP cyclohydrolase II (*ribA*) | 8 | 1.0498E-06 |
| H | *ArtHe_05135* | 6,7-dimethyl-8-ribityllumazine synthase (*ribE*) | 8 | 2.8309E-09 |
| E | *ArtHe_12855* | ABC transporter ATP-binding protein | 44 | 4.4332E-28 |
| E | *ArtHe_12860* | Extracellular solute-binding protein | 67 | 5.1686E-21 |
| E | *ArtHe_12865* | ABC transporter permease | 61 | 1.2252E-12 |
| E | *ArtHe_12870* | ABC transporter permease | 51 | 3.6566E-10 |
| M | *ArtHe_16210* | CapA family protein | 8 | 4.3305E-22 |
| P | *ArtHe_17390* | Potassium-transporting ATPase subunit (*kdpA*) | 16 | 6.6402E-15 |
| P | *ArtHe_17395* | Potassium-transporting ATPase subunit F | 134 | 5.2826E-06 |

## 4. Discussion

Extremophile microorganisms have functionalities of great interest for the biotechnological sector and, for this reason, numerous works have been oriented to isolate these microorganisms from different extremophile niches (Musilova et al., 2015; Raddadi et al., 2015; Orellana et al., 2018; Castillo et al., 2021; Tada et al., 2021). In this work, we have explored a very peculiar extreme niche developed by human technology such as solar panels. The solar panels that are currently spread over large surfaces of fields and cities on the planet mimic the changing weather conditions that can be found, for example, in deserts (Dorado-Morales et al., 2016). Therefore, they can be a source of microorganisms resistant to extreme desiccation and radiation conditions. Using a simple method, we have isolated a cultivable bacterium that shows high tolerance to desiccation and that has been identified within the genus *Arthrobacter*. The strain *Arthrobacter* sp. Helios, in contrast with *Exiguobacterium* sp. Helios, also isolated from the same solar panel (Castillo et al., 2021), has a desiccation tolerant phenotype independent of the growth phase. That is, the cells are ready to tolerate desiccation both in the exponential phase and in the stationary phase of growth. Our results show that this is not the case in a very closely related bacterium such as *A. koreensis*, which shows greater tolerance to desiccation in the stationary phase of growth like *Exiguobacterium* sp. Helios (Fig. 3). This property can be very relevant for biotechnological applications in environmental conditions, since the desiccation situations may occur at any time and it is not possible to anticipate at which metabolic state the cells can be trapped by the drought. Many bacteria prepare their metabolism in the stationary phase to resist stress situations and in particular some are capable of generating spores which makes them highly resistant to many extreme conditions (Phillips and Strauch, 2002; Jaishankar and Srivastava, 2017). However, in the case of *Arthrobacter* sp. Helios that does not produce spores, it seems to be permanently prepared to handle the desiccation stress, which makes it a very interesting bacterium for biotechnological applications. Bacteria tolerant to desiccation have been proposed not only to directly promote plant growth (Glick, 2014), but they can also protect plants against drought (Vílchez et al., 2016; Molina-Romero et al., 2017), high salinity (Vanissa et al., 2020), metals (Jan et al., 2019), organic contaminants (Zhang et al., 2014), and both bacterial and fungal pathogens (Velázquez-Becerra et al., 2013).

In addition to the desiccation tolerance, *Arthrobacter sp*. Helios shows a significant resistance to UV irradiation and a significant salinity tolerance. Both properties could be expected since the strain was isolated from a solar panel that is subjected to a high irradiation and to a high salinity due to the progressive accumulation of salts by the daily cycles of humidity and drought, moreover when these solar panels were located near the cost. However, an unexpected property was the high resistance to selenite and the formation of SeNPs. This property can be exploited for biotechnological applications since both elemental selenium and SeNPs can be used in many purposes. For example, selenium supplementation in the diet has been correlated with health benefits effect (Rayman, 2000). Moreover, SeNPs have semiconductor and photoelectrical properties and they have been used successfully in applications ranging from solar cells, photographic exposure meters, photocopiers and rectifiers (Johnson et al., 1998). SeNPs have also applications in diverse areas such as cosmetics, coatings, packaging, biotechnology and biomedicine (Thakkar et al. 2010; Karuppiah et al. 2012; Ali et al. 2013). Many bacteria have been described with the ability to produce SeNPs (Stolz et al., 2006), however the Helios strain represents an excellent candidate for SeNPs production platform due to its high level of selenite resistance, most probably due to its adaptation to the stressful condition promoted by the metal.

The analysis of the *Arthrobacter* sp. Helios genome confirmed that this bacterium is fully equipped to respond to osmotic, oxidative, UV, and heat shock stresses. Particularly, in this work, we have carried out a transcriptomic study in order to understand how *Arthrobacter* sp. Helios adapts its metabolism in response to PEG-induced water-stress that has been proposed to mimic arid stress situations (Zhao et al., 2020). Unexpectedly, only few whole omic studies have been performed to analyse the metabolic adaptations of bacteria under PEG-induced stress. Ghedira et al. (2018) studied the PEG-responding desiccome of the microsymbiont *Frankia alni* by next-generation proteomics. This nitrogen-fixing bacterium allows dicotyledonous plants to colonize soils under nitrogen deficiency and water-stress. The response of *Frankia* cells to the presence of PEG consisted of an increased in the abundance of envelope-associated proteins like ABC transporters, mechano-sensitive ion channels and CRISPR-associated (*cas*) components. Besides, unnecessary pathways, like nitrogen fixation, aerobic respiration and homologous recombination, were markedly down-regulated (Ghedira et al., 2018). Zhao et al. (2020) studied the response of the sporulating bacterium *Bacillus megaterium* FDU301 to PEG-mediated arid stress. Their main observation was the overexpression of genes related to oxidative stress and other genes related to iron uptake, sporulation stage and biosynthesis of compatible solute ectoine (Zhao et al., 2020). Biosynthesis of the compatible solute trehalose appears to be the main mechanism to face arid-stress in the α-proteobacterium *Bradyrhizobium japonicum*, a leguminous plants symbiont, although other general mechanisms such as cell membrane protection, repair of DNA damage and oxidative stress responses are involved (Cytryn et al., 2007). Using micro-array hybridizations, the group of van der Meer has studied in *Arthrobacter chlorophenolicus* A6, *Sphingomonas wittichii* RW1 and *Pseudomonas veronii* 1YdBTEX2 the response to water stress induced by addition of NaCl (solute stress) or PEG8000 (matric stress) (Johnson et al., 2011; Moreno-Forero et al., 2016). These experiments were performed testing the transcriptional response of cells transiently exposed (30 min) to a water stress rather than measuring transcription in cells growing constantly under the applied water stress (Johnson et al., 2011). Common reactions among the three strains included diminished expression of flagellar motility and increased expression of compatible solutes. In addition, a set of common genes including ABC transporters and aldehyde dehydrogenases appeared to constitute a core-conserved response to water stress. However, Gülez et al. 2012 also tested by micro-arrays the effect of PEG8000 in *Pseudomonas putida* KT2440 and observed that, in this case, PEG does not affect cell mobility and flagellar genes were not downregulated or even suggested increasing expression as the stress prolonged.

The most remarkable differences found in the *Arthrobacter* PEG6000-induced water shortage were the overexpression of different osmotic stress resistance mechanisms. Our results showed that ABC transporter genes involved in transport of iron and siderophores, as well as of other compatible solute transporters such as glycine-betaine and amino acids, were highly overexpressed compared to the control condition. This response is consistent with other studies, where bacteria exposed to different osmotic stress showed a higher expression of genes involved in the accumulation of K^+^ and different compatible solutes (Dinnbier et al., 1988; Whatmore et al., 1990; Zeidler and Müller, 2019), since it is considered a first, fast response to adjust the cell turgor against an osmotic shock (Zeidler and Müller, 2019). Surprisingly, no overexpression of genes involved in the *de novo* synthesis of compatible solutes were observed in *Arthrobacter* sp. Helios. This could be due to the high energetic cost required in their biosynthesis, being their uptake from the environment a more favourable reaction (Oren, 1999). On the other hand, we observed a 5-fold induction of *ArtHe_12305* gene, coding a mechanosensitive ion channel (MscS) family protein that becomes open in response to stretch forces in the membrane lipid bilayer, thus facilitating the efflux of all the osmolytes accumulated during the osmotic stress and avoiding a possible cell lysis (Martinac, 2001). Activation of the mechanosensitive channel McsL was found highly upregulated in the PEG induced *F. alni* proteome as well (Ghedira et al., 2018).

One of the extreme upregulated genes in PEG35 was *ArthHe_14340* (321-fold), coding a putative amino oxidase participating in the catabolism of phenylethylamine. The uptake and biosynthesis of polyamines is known to play a key role in the survival upon in response to abiotic stresses, as they participate in many cell defence strategies. Polyamines are known to play an indirect role in plant abiotic stress tolerance by participating in osmolyte synthesis in response to stress (Sengupta et al., 2015). PEG induced stress influences the plant immune response and resistance to pathogen infections by enhancing the activity of diamine oxidases and polyamine oxidases (Hatmi et al., 2014). A signal molecule such as H_2_O_2_ derived from these oxidations is known to mediate many physiological phenomena. Furthermore, the presence of polyamines stabilizes bacterial spheroplasts and protoplasts from osmotic shock (Koski and Vaara, 1991) and improve the survival rate of freeze thawed *E. coli* cells (Souzu, 1986). On the other hand, the fact that polyamines are toxic to bacteria if produced in excess might explain the high induction fold found in the PEA degradation cluster. It has been suggested that the catabolism and efflux, along with anabolism and uptake mechanism of polyamines, is vital to maintain the homeostasis in bacteria, since the accumulation of polyamines leads to inhibition of cellular growth (Karahalios et al., 1998; Banerji et al., 2021). The activity of this enzyme together with the activity of other induced lyases, such as serine ammonia-lyase, might lead to the accumulation of ammonia, that might be the reason why a high number of genes coding enzymes involved in the assimilation of nitrogen-containing compounds were downregulated. For example, ammonium and urea transporters, urease and nitrate reductase coding genes showed a significant downregulation in *Arthrobacter* sp. Helios under the studied conditions (Table 4).

*ArtHe_16370* coding a metallopeptidase is also one of the most induced genes (58-fold) in PEG35 conditions. A high induction of a metallopeptidase coding gene was also found in the *F. alni* proteome that putatively cleaves dipeptides into single peptides to provide intracellular osmolytes to offset the high extracellular osmotic pressure (Ghedira et al., 2018). *ArtHe_11205* codes a member of the Usp universal stress protein family. Usp is a small cytoplasmic bacterial protein whose expression is enhanced when cells are exposed to stress agents. Usp enhances the rate of cell survival during prolonged exposure to such conditions, and may provide a general “stress endurance” activity (Kvint et al., 2003). However, deletion of the UspE-like protein whose expression was induced in desiccation conditions in *Salmonella enterica* did not yield to any significant phenotype (Gruzdev et al., 2012), suggesting that this Usp is not directly involved in desiccation tolerance.

The comparison of the transcriptomes in PEG0, PEG10 and PGE35 shows that the response under moderate stress (PEG10) and high stress (PG35) conditions have points in common but also show significant differences. In PEG10 conditions, the cells are capable of modifying their metabolism to maintain their growth capacity almost intact, since growth differences compared to PEG0 condition were small. However, when the concentration of PEG becomes higher, the cells have to modify their metabolism to survive, limiting their growth capacity. Some common changes that can be seen between PEG10 and PEG35 are the overexpression of different potassium (*ArtHe_17390* and *ArtHe_17395*), spermidine/putrescine (*ArtHe_12855*-*ArtHe_12870*) and iron and siderophore ABC transporters, suggesting a general response to cope either with moderate or high drought stress.

One of the biggest differences observed between PEG0 and PEG10 is the overexpression of a gene cluster involved in riboflavin biosynthesis (*ArtHe_05120-ArtHe_05140*) that was no longer observed in PEG35. It is known that iron metabolism and riboflavin are intrinsically related since both are cofactors used for many enzymes involved in redox metabolism. In addition to their role as ubiquitous intracellular cofactors, flavins have recently gained attention for their involvement in a series of extracellular processes. In bacteria, iron availability influences expression of riboflavin biosynthetic genes. There is documented evidence for riboflavin involvement in surpassing iron-restrictive conditions in some species. Although the specific mechanism through which flavins increase iron reduction in this species is not addressed, it is known that bacterially secreted riboflavin acts as electron shuttle for iron reduction during extracellular electron transfer (García-Angulo, 2017; Sepúlveda-Cisternas et al., 2018).

*Arthrobacter* sp. Helios transcriptome showed a higher number of DEGs in PEG35 compared to PEG10. We have observed the overexpression of several genes encoding glycine/betaine ABC transporters (*ArtHe_04675* and *ArtHe_04665*), a universal stress protein (*ArtHe_11205*), the 8-oxoguanine deaminase (*ArtHe_13950*), the highly induced cluster involved in phenylethylamine metabolism, and a higher number of induced ABC transporters for amino acids, peptides and iron and on the other hand, we have observed the downregulation of the aquaporin encoding gene (*ArtHe_08770*). In contrast with PEG10, where few DEGs where detected and cell growth was hardly impaired, the large changes observed in gene expression in PEG35 constitutes a massive cell response to conditions where not only growth but bacterial survival starts being compromised.

Surprisingly, although *Arthrobacter* sp. Helios appears to be able to produce trehalose by different pathways we have not observed the induction of these genes, suggesting that trehalose is not involved in the adaption to matric stress in this organism.

On the other hand, we have determined that cells adapted to PEG35 are not more tolerant to desiccation than PEG0 cells (data not shown) suggesting that the adaption to matric stress is carried out by different mechanisms than the desiccation adaption and therefore, a previous adaptation to matric stress does not improve desiccation tolerance.

The results of this work allow us to anticipate that this new bacterium isolated from solar panels can be very useful for different biotechnological applications where water scarcity or drought can be a critical factor. Our current investigations are focussed on its possible utility as a plant growth promoter and as a possible biotechnological chassis for its use in contained bioreactors in low water conditions (e.g., biofilters), since preliminary experiments indicate that this bacterium can be easily modified by genetic engineering tools.

## 5. Acknowledgements

This research was supported by grants BIO2015-66960-C3-1-R, BIO2016-79736-R and PID2019-110612RB-I00 from the Ministry of Economy and Competitiveness of Spain, RTI2018-095584-B-C41-42-43-44 from the Ministry of Science and Innovation of Spain, and by Grants CSIC 2017 2 0I 015.

## Supplementary material

**Table S1**. Transcriptomic data statistics.

**Table S2**. *Arthrobacter* sp. Helios growth in the presence of different metals and metalloids. (+) Growth was detected by *OD_600_* > 0.4. (−) No growth was detected.

**Table S3**. Carbon sources used by Arthrobacter sp. Helios. (+) Growth was detected by *OD_600_* > 0.4. (−) No growth was detected.

**Table S4**. *Arthrobacter* sp. Helios genome features according to the NCBI annotation.

**Table S5**. *Arthrobacter* sp. Helios DEGs in PEG10 vs. PEG0 and PEG35 vs. PEG0 conditions according to COG database.

**Table S6.***Arthrobacter* sp. Helios DEGs in PEG10 vs. PEG0 and PEG35 vs. PEG0 conditions annotated with KEGG database.

**Table S7.** Whole *Arthrobacter* sp. Helios transcriptomic results.

**Figure S1**. Growth curve of *Arthrobacter* sp. Helios in LB medium with increasing concentrations of NaCl.

**Figure S2.** Sodium selenite reduction by *Arthrobacter* sp. Helios grown in LB medium.

**Figure S3.** Principal component analysis. Red, green and blue circles represent triplicates of *Arthrobacter* sp. Helios samples grown in PEG0, PEG10 and PEG35, respectively.

## Notes

### Competing Interest Statement

The authors have declared no competing interest.

