## Supplemental Table 7 for "Transcriptional response of the xerotolerant *Arthrobacter* sp. Helios strain to PEG-induced drought stress": Suplementary material 1.docx

**Table S1**. Transcriptomic data statistics.

| **Sample name** | **Total read bases (bp)** | **Raw reads (bp)** | **Trimmed reads (bp)** | **Q30 (%)** | **Genome mapped reads (bp)** | **Genome mapped ratio (%)** |
| --- | --- | --- | --- | --- | --- | --- |
| PEG0_1 | 8,470,090,511 | 59,256,426 | 58,328,350 | 95.52 | 57,395,096 | 98.4 |
| PEG0_2 | 8,638,092,226 | 59,904,632 | 58,951,312 | 95.49 | 57,831,237 | 98.1 |
| PEG0_3 | 8,635,326,575 | 59,457,088 | 58,642,442 | 95.68 | 57,352,308 | 97.8 |
| PEG10_1 | 8,637,411,610 | 59,713,910 | 58,909,268 | 95.63 | 57,554,354 | 97.7 |
| PEG10_2 | 8,568,619,099 | 59,255,994 | 58,419,772 | 95.49 | 57,076,117 | 97.7 |
| PEG10_3 | 8,660,621,661 | 59,922,402 | 59,014,942 | 95.37 | 57,716,613 | 97.8 |
| PEG35_1 | 8,412,146,990 | 59,743,538 | 58,880,688 | 95.97 | 52,815,977 | 89.7 |
| PEG35_2 | 8,502,848,656 | 59,384,620 | 58,295,978 | 95.52 | 52,233,196 | 89.6 |
| PEG35_3 | 8,612,624,273 | 59,736,090 | 58,793,780 | 95.68 | 53,325,958 | 90.7 |

**Table S2**. *Arthrobacter* sp. Helios growth in the presence of different metals and metalloids. (+) Growth was detected by *OD_600_* > 0.4. (-) No growth was detected.

| **Metals** | **Concentration (mM)** | **Growth** |
| --- | --- | --- |
| NiCl_2_ | 0.62 | + |
|  | 1.25 | - |
| ZnCl_2_ | 0.62 | + |
|  | 1.25 | + |
|  | 2.5 | + |
|  | 5 | + |
|  | 10 | + |
| K_2_TeO_3_ | 0.62 | + |
|  | 1.25 | + |
|  | 2.5 | + |
|  | 5 | + |
|  | 10 | - |
| NaAsO2 | 0.62 | - |
|  | 1.25 | - |
| Na_2_HAsO_4_ | 1 | - |
|  | 10 | - |
|  | 25 | - |
| CuSO_4_ | 0.4 | + |
|  | 0.8 | + |
|  | 1.6 | + |
|  | 3.1 | - |
| CdCl_2_ | 0.62 | - |
| AgNO3 | 0.62 | - |
| Pb(NO3)2 | 0.62 | + |
|  | 1.25 | + |
|  | 2.5 | - |
|  | 5 | - |
|  | 10 | - |
| Na_2_SeO_3_ | 1 | + |
|  | 5 | + |
|  | 10 | + |
|  | 50 | + |
|  | 100 | + |
|  | 150 | + |
|  | 200 | - |

**Table S3**. Carbon sources used by *Arthrobacter* sp. Helios. (+) means that growth was detected by *OD_600_* > 0.4. (-) means that no growth detected.

| **Carbon source** | **Growth** |
| --- | --- |
| Glucose | + |
| Fructose | + |
| Xylose | + |
| Succinate | - |
| Maltose | + |
| Sucrose | + |
| Galactose | - |
| Citrate | - |
| Lactose | - |
| Arabinose | - |
| Ribose | - |
| Phenol | - |
| Benzoate | - |
| 3OH Benzoate | + |
| 4OH Benzoate | + |
| Protocatechuate | + |
| Catechol | - |
| Phenylacetic acid | + |
| Terephthalate | - |
| Isophthalate | - |
| Phthalate | - |
| Pyridine | - |
| Cholesterol | + |

Tables S4, S5, S6, S7

These tables can be found in the supplementary Excel file

Figure S1

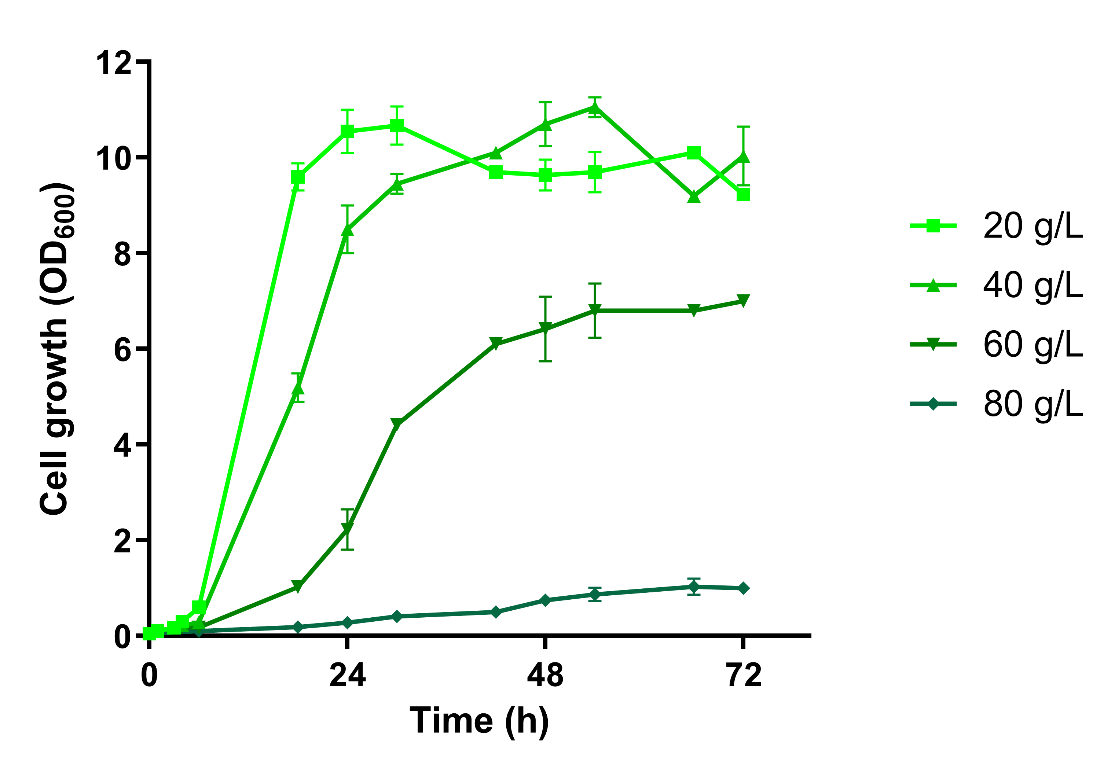

Figure S2

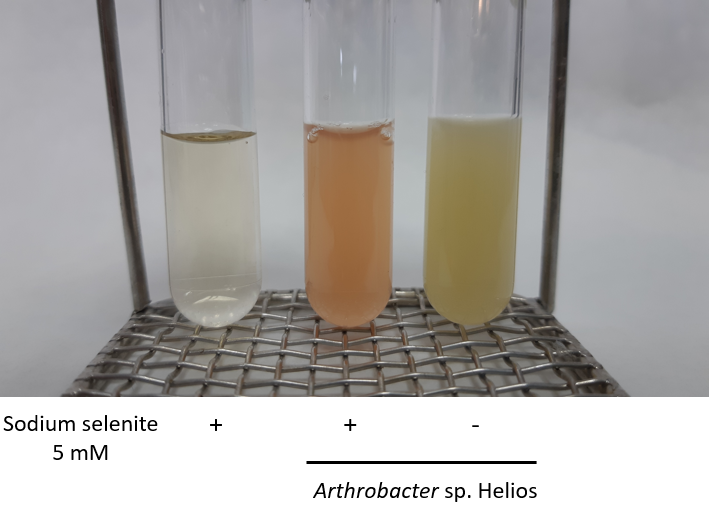

Figure S3

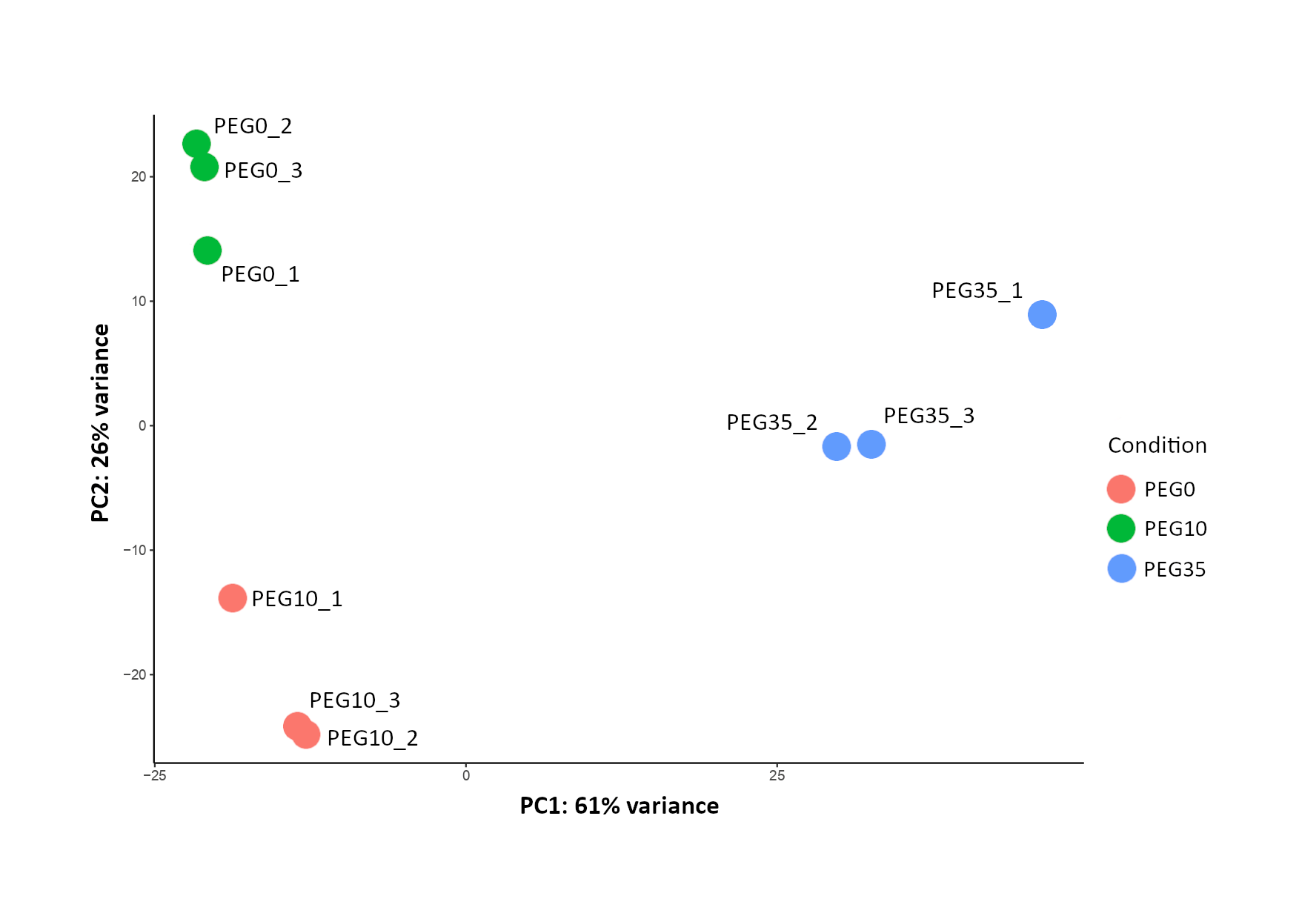
